# Spatially organized striatal neuromodulator release encodes trajectory errors

**DOI:** 10.1101/2024.08.13.607797

**Authors:** Eleanor Brown, Yihan Zi, Mai-Anh Vu, Safa Bouabid, Jack Lindsey, Chinyere Godfrey-Nwachukwu, Aaquib Attarwala, Ashok Litwin-Kumar, Brian DePasquale, Mark Howe

## Abstract

Goal-directed navigation requires animals to continuously evaluate their current direction and speed of travel relative to landmarks to discern whether they are approaching or deviating from their goal. Striatal dopamine and acetylcholine are powerful modulators of goal-directed behavior, but it is unclear whether and how neuromodulator dynamics at landmarks incorporate relative motion for effective behavioral guidance. Using optical measurements in mice, we demonstrate that cue-evoked striatal dopamine release encodes bi-directional ‘trajectory errors’ reflecting relationships between ongoing speed and direction of locomotion and visual flow relative to optimal goal trajectories. Striatum-wide micro-fiber array recordings resolved an anatomical gradient of trajectory error signaling across the anterior-posterior axis, distinct from trajectory error independent cue signals. Dynamic regression modeling revealed that positive and negative trajectory error encoding emerges early and late respectively during learning and over different time courses in the medial and lateral striatum, enabling region specific contributions to learning. Striatal acetylcholine release also encodes trajectory errors, but encoding is more spatially restricted, opposite polarity, and delayed relative to dopamine, supporting distinct roles in modulating striatal output and behavior. Dopamine trajectory error signaling and task performance were reproduced in a reinforcement learning model incorporating a conjunctive state space representation, suggesting a potential neural substrate for trajectory error generation. Our results establish region specific neuromodulator signals positioned to guide the speed and direction of locomotion to reach goals based on environmental landmarks during navigation.

## Introduction

Sensory cues experienced during goal-directed navigation serve as landmarks to direct appropriate adjustments in the direction and speed (trajectory) of ongoing movement. Studies have implicated the striatum in landmark-based navigation strategies, which complement and compete with hippocampus and cortex dependent cognitive map-based strategies^1–4^. The striatum is proposed to learn discrete associations between specific cues and actions that precede reward, providing a basis for eliciting fixed responses upon future encounters with the cue^4–7^. However, effective landmark based navigation cannot rely solely on fixed cue-action associations but requires information about whether an animals’ current trajectory, relative to a cue, is bringing it closer or further from the goal. For instance, seeing a sign for an ice cream store traveling in one direction on the highway may indicate that you’re approaching and need to continue, but seeing the same sign while traveling the opposite direction would indicate that you’re deviating and need to reverse course. In this case, the appropriate response to the cue is not fixed but depends on the relative motion of the landmark and observer, which can be calculated based on self motion and visual flow. Thus, current associative learning theories of landmark based navigation are inadequate to explain how animals can appropriately learn and adjust ongoing behavioral trajectories to reach goals. While the neural representations in the hippocampus and cortex underlying cognitive map based navigation have been widely studied, the neural signals in the striatum and interconnected regions for guiding ongoing behavioral trajectories relative to landmarks are unresolved.

Striatal neuromodulator release is widely appreciated to contribute to cue-action learning by reinforcing associations and motivating actions associated with reward^8–12^. In prominent reinforcement learning models, dopamine release to cues, actions, and rewards encodes a reward prediction error (RPE), which promotes the formation of cue-action associations to drive future choice^8,11,13^. Acetylcholine release from cholinergic interneurons also partially reflects RPEs, but with an opposite polarity to dopamine^14–16^ and may counteract the influence of dopamine to promote behavioral extinction and flexibility^17–20^. RPEs to cues and actions scale with expected reward probability, magnitude, and timing, allowing for regulation of learning and motivation based on predicted costs and benefits^21–24^. Despite these insights, an understanding of how striatal neuromodulator release to cues integrates continuous, relative motion of the cue and observer, information necessary for landmark-based navigation, is lacking. As illustrated in the example above, an RPE signal which indicates only that a road sign is associated with ice cream would be ineffective for guiding accurate future or current navigation because it does not incorporate the direction and speed of movement relative to the sign. Therefore, in addition to cue identity indicating the likelihood of future reward, cues serving as landmarks must also elicit neural signals to direct appropriate adjustments in the direction and speed (trajectory) of ongoing relative movement during navigation.

Dopamine and acetylcholine signaling have primarily been measured in instrumental tasks in which animals are motionless at cue presentation or Pavlovian tasks in which motion of the cue and animal is irrelevant to obtaining reward. Several studies have reported ramping of striatal dopamine release as animals approach goals during navigation, a signal believed to encode goal proximity, perhaps via RPEs, and regulate motivation based on continuously changing sensory cues^25–29^. However, in these tasks, the direction and speed of motion relative to the environment was fixed along a learned approach trajectory, so animals were always moving towards the goal. Thus, previous experimental designs have not permitted investigation of whether neuromodulator signals reflect cue and animal motion relative to goals. We addressed this gap by optically measuring striatum-wide dopamine and acetylcholine release using high density micro-fiber arrays^30^ in head-fixed mice performing visually guided tasks which simulate features of landmark based navigation. Our study reveals ‘trajectory error’ signals in cue-evoked neuromodulator release, reflecting the direction and speed of locomotion and cues relative to goals, based on instructional cue identity. Our large scale measurements resolved temporal and anatomical gradients in the magnitude and learning time course of dopamine and acetylcholine trajectory error signaling across striatal regions. These signals contain the information necessary to guide the ongoing trajectory of goal-directed navigation relative to specific external landmarks in real time or across learning.

## Results

### Measurements of striatum-wide dopamine release with micro-fiber arrays during a visually guided instrumental task

We designed a task capturing key features of landmark based navigation, in which animals experienced cues while in continuous motion and were required to make appropriate adjustments based on the cue identity. Mice were head-fixed with their limbs resting on a floating ball, allowing them to locomote in 2-dimensions^31^. On each trial, they were presented with one of two visual cues at the center of an array of computer monitors (Fig. 1a). The angular velocity of the mouse movement on the ball was translated into movement of the visual cue across the screens. To receive a water reward or end the trial, the mice were required to move the cue laterally across the screens by ∼33 degrees in either direction. Trials lasted for a median 5.0 s from cue onset to reward delivery, with an average of 111 trials for each 33 minute recording session. Each cue was presented pseudorandomly on 50% of trials and the cue pattern indicated which direction was rewarded. Mice (n = 10) were trained to perform with a reward rate of over 80% for at least 3 consecutive days, indicating stable task performance (Fig. 1b). Importantly, the mice ran freely with a range of angular velocities before cue onset (Fig. 1b). On each trial, the cue identity indicated to the mouse whether they were currently traveling in a direction towards (congruent) or away (incongruent) from the rewarded direction signaled by the cue. For congruent trials during correct task performance, mice continued to run in the same direction, while on incongruent trials they switched direction (Fig. 1c,d). Thus, unlike standard instrumental tasks, mice did not initiate new actions from rest but adjusted their ongoing locomotion patterns in accordance with the cue, a situation analogous to encountering changing visual landmarks during ongoing navigation.

**Figure 1:**
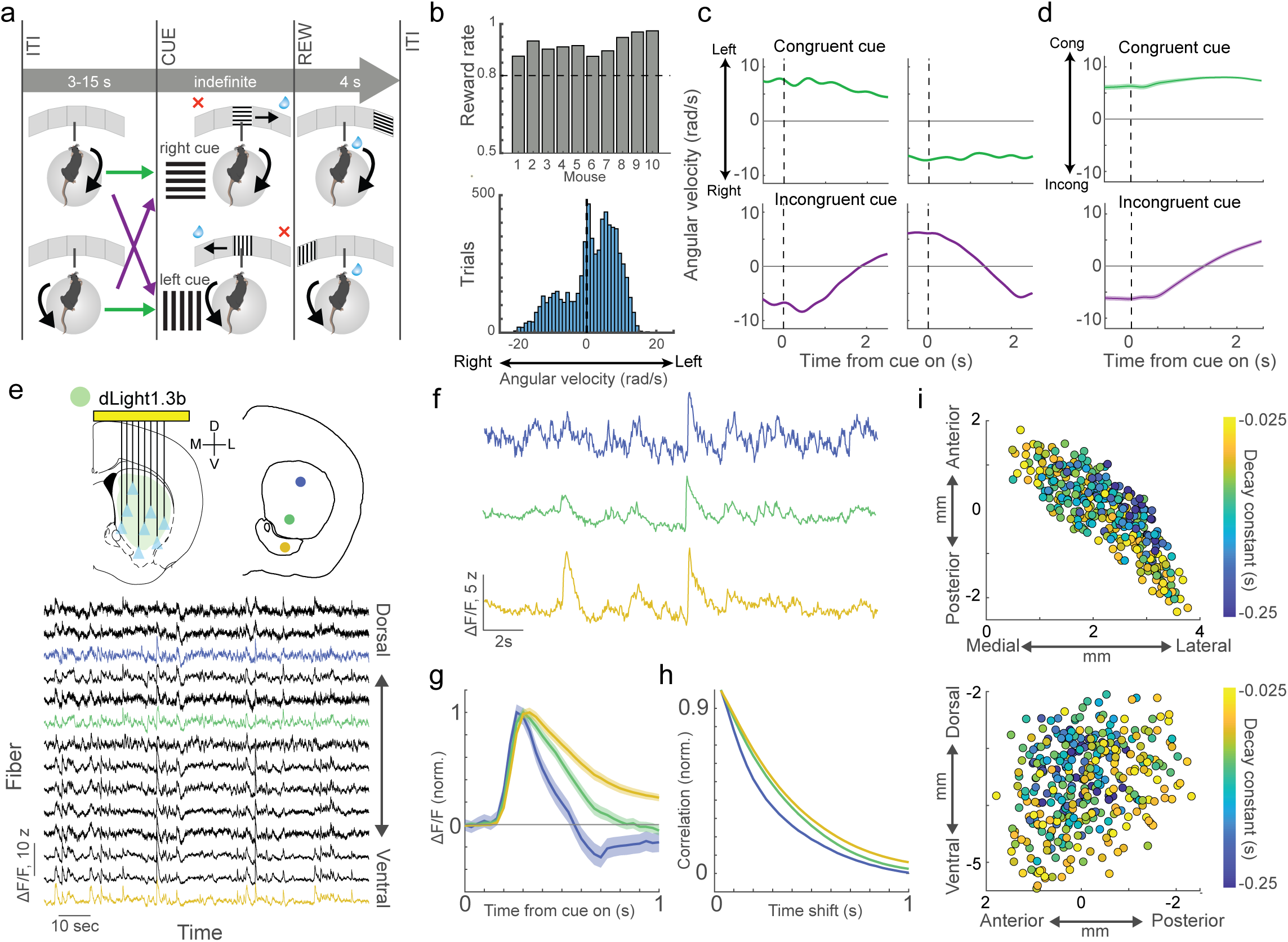
Temporal organization of striatum-wide dopamine release during a visually guided instrumental task. **a,** Instrumental task schematic. Mice ran on a spherical treadmill during the ITI with angular velocities that were either congruent (green arrow) or incongruent (purple arrow) with the rewarded direction indicated by a visual cue presented on each trial. At cue onset, treadmill movement was converted into lateral movement of the visual cue across the screens and reward delivery occurred when the cue reached a fixed lateral position. **b,** Top, average reward rate across post-learning sessions for individual mice. Bottom, histogram of angular velocities for each trial averaged within a 300ms window before cue onset. Positive and negative values indicate angular velocity in the left and right directions respectively. **c,** Mouse angular velocity, signed by running direction (+ left, - right) on four example trials aligned to cue onset where the mouse was running to the left or right in the congruent (top, green) or incongruent (bottom, purple) directions prior to cue onset. **d,** Mean congruence-signed angular velocity aligned to cue onset for all congruent (top) and incongruent (bottom) trials for a single mouse (n = 321 and 297 trials respectively). The angular velocity is signed relative to the congruent (+) or incongruent (-) direction, regardless of left or right running direction. **e,** Top left, schematic of striatum-wide dopamine release measurements with micro-fiber arrays. Bottom, example z-scored ΔF/F traces recorded simultaneously from 14 different fibers in a single animal, ordered from dorsal (top) to ventral (bottom). Fiber locations for the three colored traces are indicated in the coronal section on the top right. **f,** Example ΔF/F traces with different temporal dynamics. Colors correspond to the traces in e. **g,** Trial-averaged, peak-normalized ΔF/F aligned to cue onset for the same three fibers in f (n = 762 trials). **h,** Autocorrelograms calculated across the entire recording period normalized to the maximum correlations for the three fibers in f. **i,** Horizontal (top) and sagittal (bottom) maps of autocorrelation decay time constants of ΔF/F for each fiber (circle) across all mice (n = 351 total fibers in 10 mice). Locations are relative to bregma. Shaded regions in all plots are 95% confidence intervals.

Encoding of reward prediction errors and other behavioral variables in dopamine release is known to vary across striatal regions^32^, so we applied a multi-fiber array approach^30^ to optically measure dopamine release at many locations simultaneously (24-57 fibers/mouse, 351 total fibers) across the 3-dimensional striatum volume during task performance (Fig. 1e, Extended Data Fig. 1). This approach yielded high signal to noise measurements of rapid dopamine release using the fluorescent sensor dLight1.3b^33^, which could be isolated from hemodynamic and motion artifacts using near isosbestic illumination (Methods). Across all task time, we found spatial gradients in the intrinsic timescale of dopamine signaling, with the most rapidly decaying signals in the dorsolateral striatum and slower decaying signals in the ventromedial and posterior striatum (Fig. 1f-i). These results extend previous studies which have reported differences in dopamine release kinetics and frequency across the dorsal-ventral axis during behavior^34,35^.

### Cue-evoked dopamine release encodes spatially organized errors in locomotion direction

Dopamine release transiently increased at cue onset for both congruent and incongruent trials, but release on congruent trials was significantly higher than incongruent at most recording sites (Fig. 2a-c, 290/351 congruent higher, 4/351 incongruent higher, one-tailed shuffle test, p<0.01, n = 10 mice). For sessions with isosbestic illumination, only a small fraction of fibers had significant differences between trial types for 405 nm illumination with no congruent preference (Fig. 2a,b; 27/246 congruent higher, 33/246 incongruent higher; 185/246 cong. higher for quasi-simultaneous 470 nm illumination, one-tailed shuffle test, p<0.01, n = 7 mice). At many locations (178/351 fibers in 10 mice), the cue-evoked release significantly dipped below baseline on incongruent trials (Extended Data Fig. 2a-c, triggered average below the 99% confidence interval of a null distribution for at least 3 timepoints in a row). Thus, cue-evoked striatal dopamine release encodes positive (on congruent trials) and negative (on incongruent trials) errors between locomotion direction and the rewarded direction. Differences in cue-evoked dopamine between congruent and incongruent trials were present regardless of the cue identity or locomotion direction (ipsi- or contralateral relative to the implant), though more locations were significant for contralateral running trials (Extended Data Fig. 2d). Thus, unlike related dopamine reward prediction errors described previously^24,36,37^, these signals were not purely reflective of fixed, learned reward probabilities associated with particular actions or cues (as correct performance was high for all cues and directions) but indicated whether the animals’ direction of motion, relative to the cue identity, was bringing it closer or further from the goal.

**Figure 2:**
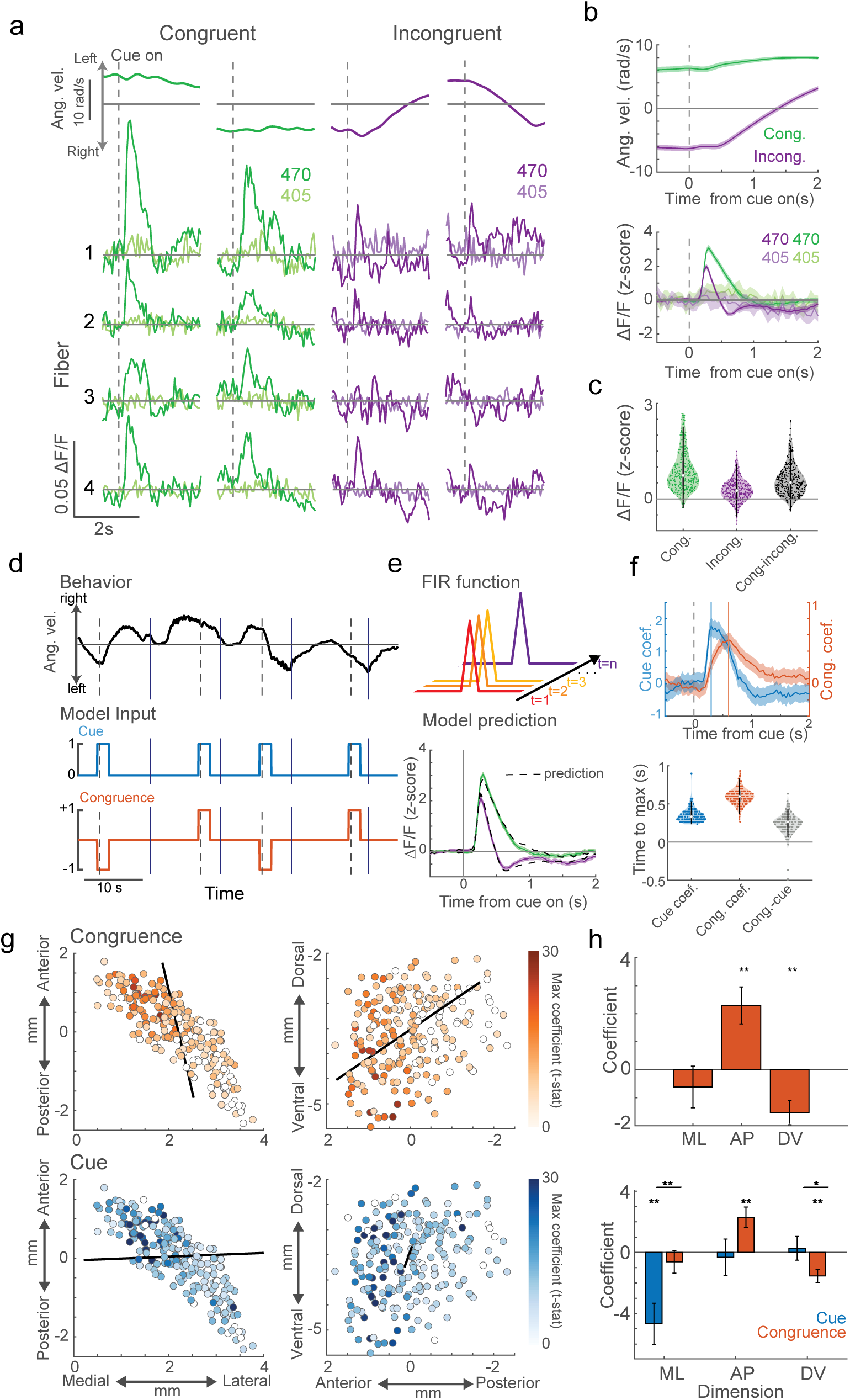
Dopamine release at cue onset encodes the congruence between the cue identity and ongoing movement direction. **a,** Mouse angular velocity, signed by running direction (+ left, - right) (top) and ΔF/F (bottom) for four example fibers aligned to cue onset on two congruent (green) and incongruent (purple) trials. Dark colors, 470nm illumination, light colors, 405nm. **b,** Top, congruence-signed angular velocity (+ congruent, - incongruent) aligned to cue onset averaged across all rewarded congruent (green) and incongruent (purple) trials for one example mouse. Bottom, average ΔF/F for the same trials at top for a single representative fiber. (n = 321 congruent 470 nm trials; n = 297 incongruent 470nm trials; n = 44 congruent 405nm trials; n = 39 incongruent 405nm trials). **c,** Violin plots of the average ΔF/F from 0.5-1 s after cue onset across fibers and mice for congruent trials (green), incongruent trials (purple) and the difference (congruent-incongruent, grey) (n = 351 fibers, 10 mice). **d,** Model inputs for cue (blue) and congruence (orange) finite impulse response (FIR) functions for an example task period (angular velocity, black; cue onset, dashed lines; reward, solid lines). **e,** Top, schematic of a standard FIR function. Bottom, average ΔF/F aligned to cue onset for congruent (green) and incongruent (purple) trials, with the model prediction for each trial type overlaid (dashed line). **f,** Top, model coefficients for cue (blue) and congruence (orange) terms at each time point relative to cue onset for a single fiber. Dashed lines show the time of the maximum coefficient. Bottom, violin plots across all fibers and mice of the times to maximum coefficient, relative to cue onset for cue (blue) and congruence (orange) coefficients, and the difference between congruence and cue (grey). **g,** Horizontal (left) and sagittal (right) maps of t-statistics for the maximum congruence (top, orange color scale) and cue (bottom, blue color scale) coefficients for each fiber (circle) across all mice. Black line indicates the axis of maximal variation (Methods). Empty circles, not significant (t-test on model coefficients, p<0.05 for 3 timepoints in a row within 1s cue window, Bonferroni corrected). **h,** Coefficients of a spatial gradient analysis, indicating the variation in coefficient t-stats along medial-lateral (ML), anterior-posterior (AP), and dorsal-ventral (DV) axes for the congruence (top) and congruence and cue comparison (bottom, orange and blue respectively). Error bars, standard error (t-test on the model coefficients and two-way ANOVA for the interactions, *p<0.05, **p<0.01). Shaded regions in all plots are 95% confidence intervals.

To isolate the effect of the cue-direction congruence on cue-evoked dopamine release, we modeled release at each timepoint after cue onset as a linear combination of finite impulse response functions (Methods, Fig. 2d,e). This analysis revealed two temporally and spatially distinct components of the dopamine signals: an early dopamine rise that was independent from the cue-direction congruence (mean latency 0.35s), and a later ‘congruence sensitive’ component (mean latency 0.60s, Fig. 2f). Positive congruence encoding reflected higher dopamine release on trials where the direction of travel at cue onset was correct, relative to incorrect. Although congruence encoding was present across most striatum locations, there was a clear spatial gradient across the striatum (174/213 significant fibers from 6 mice, t-test on model coefficients, p<0.05 for 3 timepoints in a row within 1s cue window, Bonferroni corrected): congruence encoding decreased from anterior-ventral to posterior-dorsal (Fig. 2g,h t-test on spatial coefficients, ML: p = 0.407, AP: p = 6e-4, DV: p = 4e-4). Interestingly, while congruence encoding was present across the dorsal-ventral extent of the anterior striatum, the largest dips below baseline on incongruent trials were concentrated in the anterior dorsolateral striatum (Extended Data Fig. 2a-c), perhaps due to the faster temporal dynamics in that region (Fig. 1i, see Discussion). Gradients in congruence encoding were significantly different in the ML and DV dimensions relative to the earlier congruence-independent (‘cue’) response, which displayed a medial to lateral gradient (Fig. 2g,h, two-way ANOVA on the interactions, ML: p = 0.0088, AP: p = 0.057, DV: p = 0.042). These results suggest that congruence encoding represents a distinct computation, which is separated in time and space from a congruence-independent response reflecting velocity-independent properties of the cue.

### Dopamine congruence signaling scales with locomotor speed at cue onset

Mice ran with a range of locomotor speeds on the treadmill prior to cue onset (Figs. 1b and 3a,b), so we next tested whether the congruence encoding scaled with speed. This scaling represents *how* right or wrong an animals’ ongoing locomotion vector was at cue onset. We modified the binary direction/cue identity congruence term in the model to scale with angular speed at cue onset (Fig. 3d), and refer to the new combined congruence-speed term as the ‘trajectory error’ (TE). At many locations, the cue-evoked dopamine signals scaled with TE, with the largest average signals for the fastest congruent trials and the smallest for the fastest incongruent (Fig. 3c-e, 241/351 fibers significant across 10 mice, t-test on model coefficients, p<0.05 for 3 timepoints in a row within 1s cue window, Bonferroni corrected). This scaling was not present at any recording site during 405 nm illumination (Extended Data Fig. 3, 0/246 fibers significant across 7 mice, 103/246 significant for quasi-simultaneous 470 nm illumination). Further, scaling was continuous across TE magnitudes and the spatial patterns were not affected by differences in pre-cue baseline ΔF/F on congruent or incongruent trials (Extended Data Fig. 4). Replacing the congruence term with a scaling TE term improved the model fit at ∼70% of recording sites (Fig. 3f, 148/213 sites in 6 mice with lower AIC for TE model), and the spatial patterns and timing of TE encoding overlapped strongly with congruence encoding, indicating that congruence encoding predominantly scales with speed (Fig. 3e,g-i). Most fibers (48%) had significant TE encoding for both congruent and incongruent trial types when considered independently, and a smaller fraction of sites scaled significantly with TE only on congruent (19%) or incongruent (15%) trials (Extended Data Fig. 5). Spatial patterns for congruent and incongruent trials were similar, but notably congruent TE encoding was present across the anterior-posterior axis, while incongruent TE encoding was localized to the anterior striatum, with a dorsal bias (Extended Data Fig. 5). Together these results illustrate that spatially organized, cue-evoked dopamine release encodes positive and negative trajectory errors which scale with a combination of continuous velocity magnitude and direction congruence with the cue identity.

**Figure 3:**
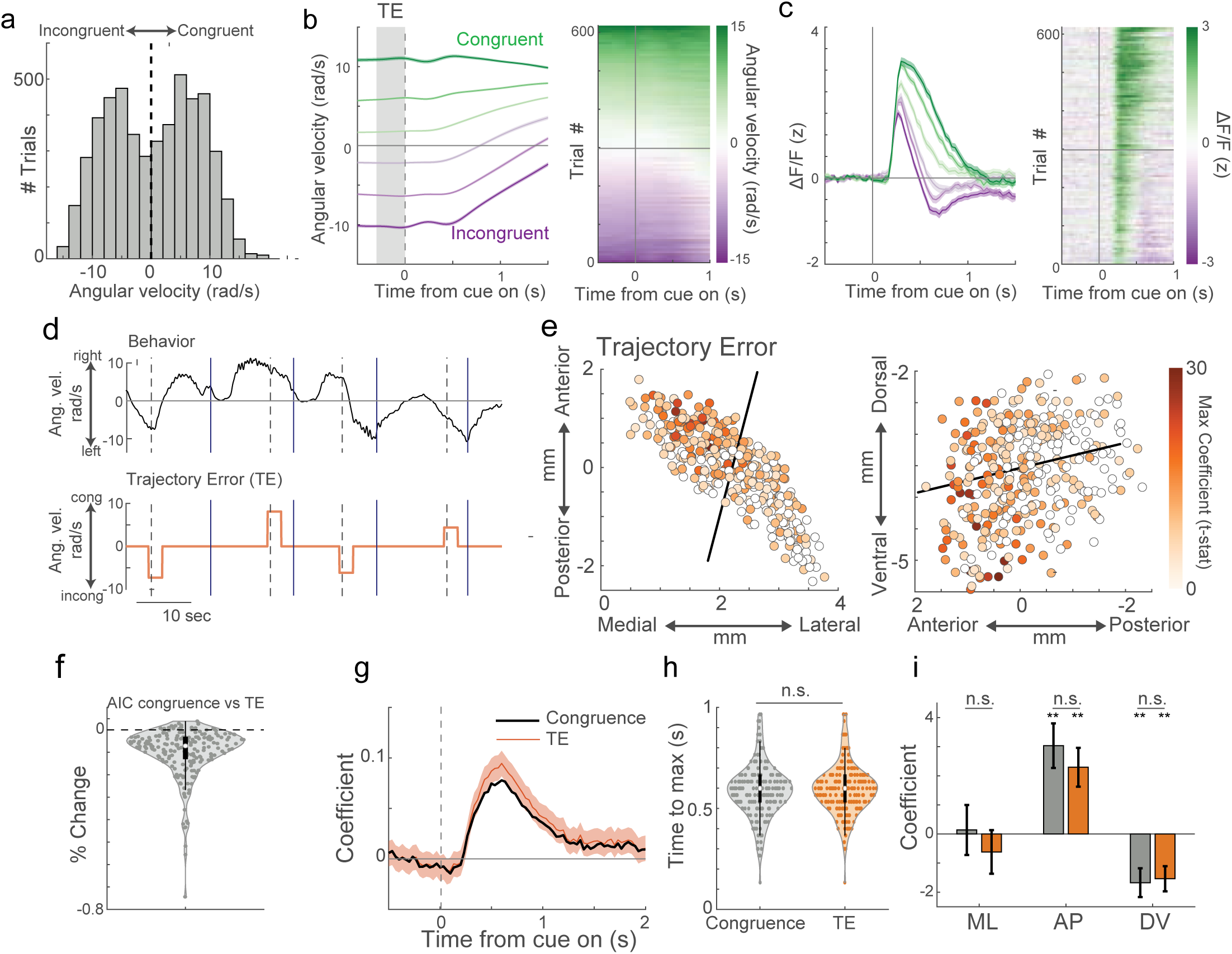
Spatially organized dopamine release encodes a trajectory error reflecting a conjunction of continuous angular velocity and direction relative to the cue identity. **a,** Histogram of congruence-signed angular velocities (+ congruent, - incongruent) prior to cue onset for each trial across all trials and mice (n = 5066 trials across 10 mice). **b,** Left, average congruence-signed angular velocity aligned to cue onset for congruent (green) and incongruent (purple) trials split into thirds by magnitude (dark to light colors indicates fast to slow) for one mouse. Grey region indicates time over which trajectory error was calculated. Shaded regions, mean +/- S.E.M. Right, raster plot of the congruence-signed angular velocity on for all trials included on left aligned to cue onset, sorted by TE. Horizontal line indicates the TE closest to 0. **c,** Same as b, but for ΔF/F for one representative fiber in the same mouse as b. Rasters are sorted by TE magnitude as in b, right. **d,** Angular velocity (black) and schematic of model inputs for the trajectory error model (orange) for an example task period. The trajectory error FIR functions tile the cue period and scale with the magnitude of the trajectory error on each trial. Cue onset, dashed lines; reward, solid lines. **e,** Horizontal (left) and sagittal (right) maps of t-statistics for maximum trajectory error coefficients for each fiber (circle). Black line indicates the axis of maximal variation. Empty circles, not significant (t-test on model coefficients, p<0.05 for 3 timepoints in a row within 1s cue window, Bonferroni corrected). **f,** Violin plot of the percent change in Akaike information criterion (AIC) from the congruence model to the trajectory error model across all fibers and mice. Improved model fit indicated by negative AIC changes. **g,** Model coefficients for the trajectory error (orange) and congruence (black) models for an example fiber. Shaded region, 95% confidence interval. **h,** Violin plots of the times to max coefficient peak relative to cue onset for congruence (grey) and trajectory error (orange) models for each fiber. (n.s., not significant, two-tailed shuffle test, p = 0.4342, n = 174 fibers significant for congruence, n = 179 fibers significant for trajectory error). **i,** Spatial coefficients for the congruence (grey) and trajectory error (orange). Error bars, mean +/- S.E.M. (t-test on model coefficients, ANOVA on interaction *p<0.05, **p<0.01, n.s. not significant).

### Trajectory error encoding is not explained by task independent locomotion relationships

Evidence has emerged that dopamine neuron activity and region-specific striatal dopamine release encode aspects of locomotion, including acceleration magnitude and direction^38–45^. Given that TEs scale with locomotion speed prior to cue onset and relate inherently to movement adjustments to the cue, we addressed the possibility that TE encoding in dopamine release may be impacted, or even solely explained by, locomotion parameters per se (Extended Data Fig. 6). A model including trial-by-trial acceleration terms was a better fit than a model without the acceleration terms for ∼75% of recording locations, indicating that locomotor acceleration contributes to dopamine signaling and could, at least partially, be captured by our model (Extended Data Fig. 6a, 267/351 fibers in 10 mice with lower AIC with acceleration terms). However, TE coefficient magnitudes were unaffected by removing acceleration terms from the model at the majority of recording sites (Extended Data Fig. 6b,c, 5/351 fibers in 10 mice with a significant difference in peak TE, t-test, p<0.05), and spatial patterns of the dopamine TE encoding were not significantly affected by the acceleration terms (Extended Data Fig. 6d, ANOVA on interactions, ML: p = 0.85, AP: p = 0.73, DV: p = 0.92). These results indicate that TE encoding is largely independent from acceleration encoding.

To further investigate whether movement per se may account for TE signaling, we examined the relationship between dopamine release and angular acceleration during the intertrial interval. To determine whether correlations with acceleration are dependent on running direction, we considered contralateral and ipsilateral (relative to the implant side) running periods separately and correlated release with the derivative of speed (absolute value of velocity) in each direction. Release magnitude was significantly positively correlated with acceleration during contralateral running and negatively correlated during ipsilateral running at 94/213 sites (Extended Data Fig. 6e,f, t-test on model coefficients, p<0.01, n = 6 mice), indicating that dopamine increases for contralateral accelerations (and/or decreases for decelerations) and vice versa for ipsilateral accelerations. Correlations were strongest across a restricted region of the dorsolateral striatum, consistent with the input topography of movement related dopaminergic cell types and previous reports of dopamine signaling in this region^39,44,46^. Thus, dopamine encodes angular acceleration with respect to direction, with regional specificity that partially overlaps with the anatomical distribution of TE encoding. Angular acceleration was oppositely correlated with TE on contralateral and ipsilateral trials (Extended Data Fig. 6g,h). Therefore, if acceleration per se were accounting for dopamine TE scaling, TE encoding should be oppositely signed on contralateral and ipsilateral trials. Contrary to this, TE encoding was positive for both trial types, and the spatial patterns were not significantly different (Extended Data Fig. 6i-k, ANOVA on interactions, ML: p = 0.68, AP: p = 0.51, DV: p = 0.64). These results establish a restricted spatial organization of striatum-wide encoding of spontaneous directional acceleration and provide further support that TE encoding is not driven by or significantly affected by acceleration encoding per se.

Finally, we tested whether cue-evoked dopamine release scaled with mouse velocity at cue onset, independently of the task, by measuring dopamine release to simple Pavlovian cues which did not signal the reward direction (Extended Data Fig. 7). There was no significant encoding of velocity magnitude at the time of cue onset at any recording site (0/150 fibers significant, n = 4 mice), confirming that TE encoding reflects task relevant information about the relationship between the instructional cue and ongoing directional locomotion rather than spontaneous velocity per se.

### Dopamine trajectory errors can be relative to both locomotion and visual feedback

In the instrumental task (Fig. 1a), angular treadmill velocity was directly linked to visual feedback through movement of the cue. Thus, dopamine TE signals could be computed based on the direction and speed of either the locomotion, visual flow, or a combination of both. To determine whether TE encoding could be computed based only on visual cue movement, we trained naive mice (n = 4) on a passive, Pavlovian version of the task (vis-only). In the vis-only task, the stimuli were identical to the original (loc + vis) task, but mice had no control over the cue movement (Fig. 4a). Cue speed and direction were yoked to selected trials of the loc + vis task performed by separate mice to ensure an even mix of congruent/incongruent and correct/incorrect trials in each direction (see Methods). After only a few days, mice pre-licked more for correct than incorrect trials, indicating successful learning of the cue-direction association (Fig. 4b,c). There was no relationship between the visual TE calculated from the initial directional movement of the cue and a locomotion TE based on the angular velocity at cue onset (Fig. 4d-g and Extended Data Fig. 8a,b). We modeled the cue-evoked dopamine release as in the loc + vis task (Fig. 3) to test whether the signals scaled with the locomotion-independent visual TE. We found significant visual TE encoding at the majority of recording locations, with no significant effect of the task irrelevant locomotion-based TE at any recording location (Fig. 4g, 113/124 fibers significant for cue TE, 0/124 for loc TE, in 4 mice, t-test on model coefficients, p<0.05 for 3 timepoints in a row within 1.5s cue window, Bonferroni corrected). The spatial distribution of visual TE encoding across the striatum was similar to the instrumental loc+vis task, with the strongest signaling concentrated in the anterior ventral striatum (Extended Data Fig. 8c). Importantly, TE encoding was significant when computed separately for congruent or incongruent trials at a majority of recording sites (Extended Data Fig. 8e,f; 105/124 significant for both or either; 65/124 significant for both, 35/124 for congruent only, 5/124 for incongruent only in 4 mice), further confirming that dopamine cue responses scale with visual TE magnitude, rather than just encoding congruence. Together, these results indicate that cue-evoked dopamine release can reflect the congruence between the cue identity and the direction and speed of visual flow with respect to goals, independently of ongoing locomotion.

**Figure 4:**
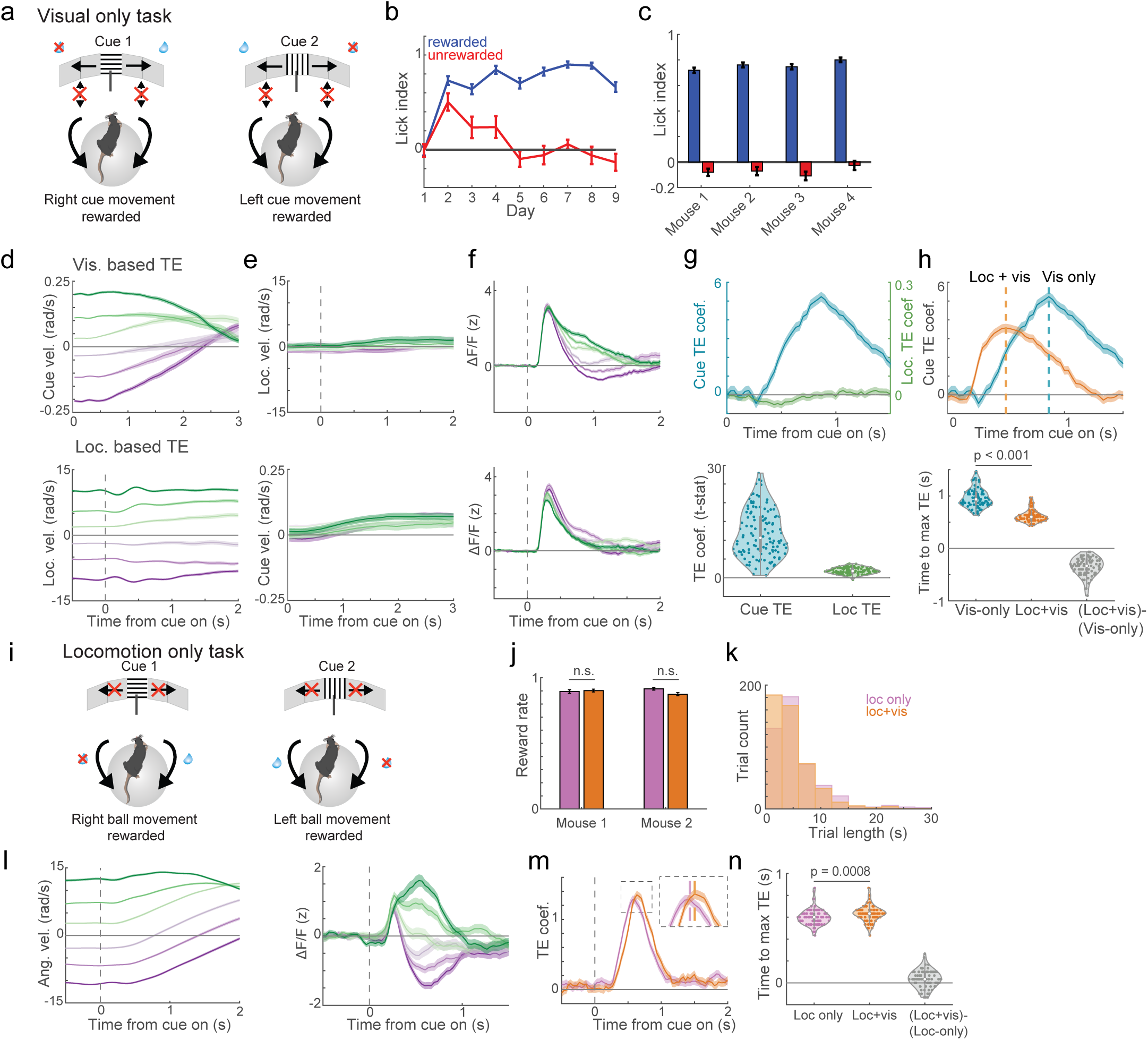
Trajectory error encoding can be based independently on visual flow and locomotion. **a,** Schematic of the visual only task (vis-only) in which the associations between cue identity and rewarded location were the same as in the instrumental (loc + vis task, Figure 1), but the mouse angular velocity was decoupled from movement of the visual cue. Direction and velocity of the cue was yoked to congruent and incongruent trials performed by separate mice (Methods). **b,** Average lick index (Methods) for rewarded (blue) and unrewarded (red) trials for each training day for one example mouse showing successful discrimination of cue-direction associations after learning. **c,** Average lick index across trials and days for rewarded (blue) and unrewarded (red) trials for each mouse. **d,** Top, average congruence-signed cue angular velocity (+ congruent, - incongruent) aligned to cue onset on congruent (green) and incongruent (purple) trials, split into thirds by cue velocity magnitude for one example mouse. Bottom, average mouse angular velocity signed by theoretical congruence as in the instrumental task, for congruent (green), and incongruent (purple) trials, split into thirds by mouse velocity magnitude. **e,** Average mouse (top) and cue (bottom) congruence-signed angular velocity for the groups of trials in d showing decoupling of cue and mouse velocity in the vis-only task. **f,** Average cue aligned ΔF/F for an example fiber for congruent (green) and incongruent (purple) trials based on the cue (top) or mouse (bottom) congruence-signed angular velocity, split into thirds by magnitude, as in d. **g,** Top, Trajectory error coefficients aligned to cue onset based on the congruence-signed angular velocity of the cue (blue) and the mouse locomotion (green) for one example fiber. Bottom, Violin plot of the max cue and locomotion based TE coefficients (t-stats) for all fibers and mice (n = 124 fibers, 4 mice). **h,** Top, trajectory error coefficients aligned to cue onset based on congruence-signed angular velocity of the cue in the vis-only task (blue) and the loc + vis task (orange) for one example fiber. Bottom, violin plot of the average timing of the max TE coefficient in the vis-only and loc+vis tasks for significant fibers and mice. (n = 91 fibers, 4 mice; two-tailed shuffle test, p<0.001). **i,** Schematic of the locomotion only task (loc-only) in which task rules were the same as loc+vis but the visual cue remained stationary at the center of the screens. **j,** Average reward rates in the loc-only (pink) and loc + vis tasks (orange), for sessions above criterion (mouse 1: n = 535 trials loc-only, 762 trials loc + vis, p = 0.23, mouse 2: n = 786 trials loc-only, 792 trials loc + vis, p = 0.10, chi-squared test). **k,** Histogram of times to reward for each task. **l,** Average congruence-signed mouse angular velocity (left) and ΔF/F for a representative fiber (right) aligned to cue onset for congruent (green) and incongruent (purple) trials, split into thirds by magnitude, during the loc-only task. **m,** Trajectory error coefficients for the loc-only (pink) and loc + vis tasks (orange) for one representative fiber. Colored dashed lines in the inset show the timing of the peak for each. **n,** Violin plots showing the timing of the maximum trajectory error coefficients for the loc-only (pink) and loc + vis (orange) task for all fibers. (n = 82 fibers, 2 mice, two-tailed shuffle test, p = 0.0008). Error bars and shaded regions in all plots are mean +/- S.E.M.

We then tested whether dopamine TE encoding could also be computed based purely on the congruence between the cue and locomotion speed and direction without visual feedback. After training on the loc+vis task, mice (n = 2) were trained on a new task (loc-only) in which cue-direction contingencies were identical to the loc + vis task, but after trial onset, the instructional cue remained stationary at the center of the visual field (Fig. 4i). As in the loc+vis task, mice ran with a range of angular velocities prior to cue onset (Fig. 4l), and they successfully adjusted their locomotion according to the cue-direction congruence, with no significant difference in accuracy compared to the loc+vis task (Fig. 4j, chi-squared test p = 0.2427 for mouse 1 and p = 0.104 for mouse 2). Significant TE encoding in cue-evoked dopamine release was present at the vast majority of recording locations in the loc-only task (Fig. 4l and Extended Data Fig. 8d, 83/85 fibers significant, t-test on model coefficients, p<0.05 for 3 timepoints in a row within 1s cue window, Bonferroni corrected). The spatial distribution of TE signaling in the loc only task was similar to the loc + vis (Extended Data Fig. 8d). Thus, TE encoding in dopamine release can be computed based on the congruence between the cue identity and the direction and speed of locomotor velocity, independently of visual feedback. Interestingly, while peak TE encoding in the vis-only task was slower than in the loc + vis task, TE encoding during the loc-only task was faster than the loc + vis task at the majority of recording sites and at the population level (Fig. 4m,n, 51/82 sites faster for loc-only, 91/91 sites slower for vis-only). Thus, TE encoding during the loc + vis task may represent a sum of distinct fast and slow computations based on locomotor and visual input respectively.

### Trajectory error encoding evolves differently across the striatum during learning

To gain insight into how dopamine TEs evolve, we recorded dopamine release across learning in a subset of mice (n = 3). A dynamic regression model^47^ was used (see Methods) to estimate the evolution of TE encoding on single trials across learning. Across recording sites and mice, TE coefficients increased rapidly and reached statistical significance well before performance exceeded chance levels (Fig. 5a,b and Extended Data Fig. 9a; Mean lag between TE and performance, 395 trials; S.E.M. 5 trials). These results indicate that dopamine TE signals may contribute to learning and further support the notion that TEs do not result from movement encoding per se. Although early emergence of TE encoding was prevalent across locations, the time course of the encoding differed across striatal axes. Nonnegative matrix factorization was used to identify changes in TE encoding early and late in learning (Fig. 5c, Methods). This revealed a bias towards early changes in TE encoding in the anterior medial striatum and a bias towards later changes in TE encoding in the central lateral striatum (Fig. 5c,d; Extended Data Fig. 9b,c), a spatial organization that was consistent across mice (Extended Data Fig 9b,c). These results indicate that TE encoding emerges along different time courses across the striatum, perhaps reflecting participation in different phases of learning or task performance.

**Figure 5:**
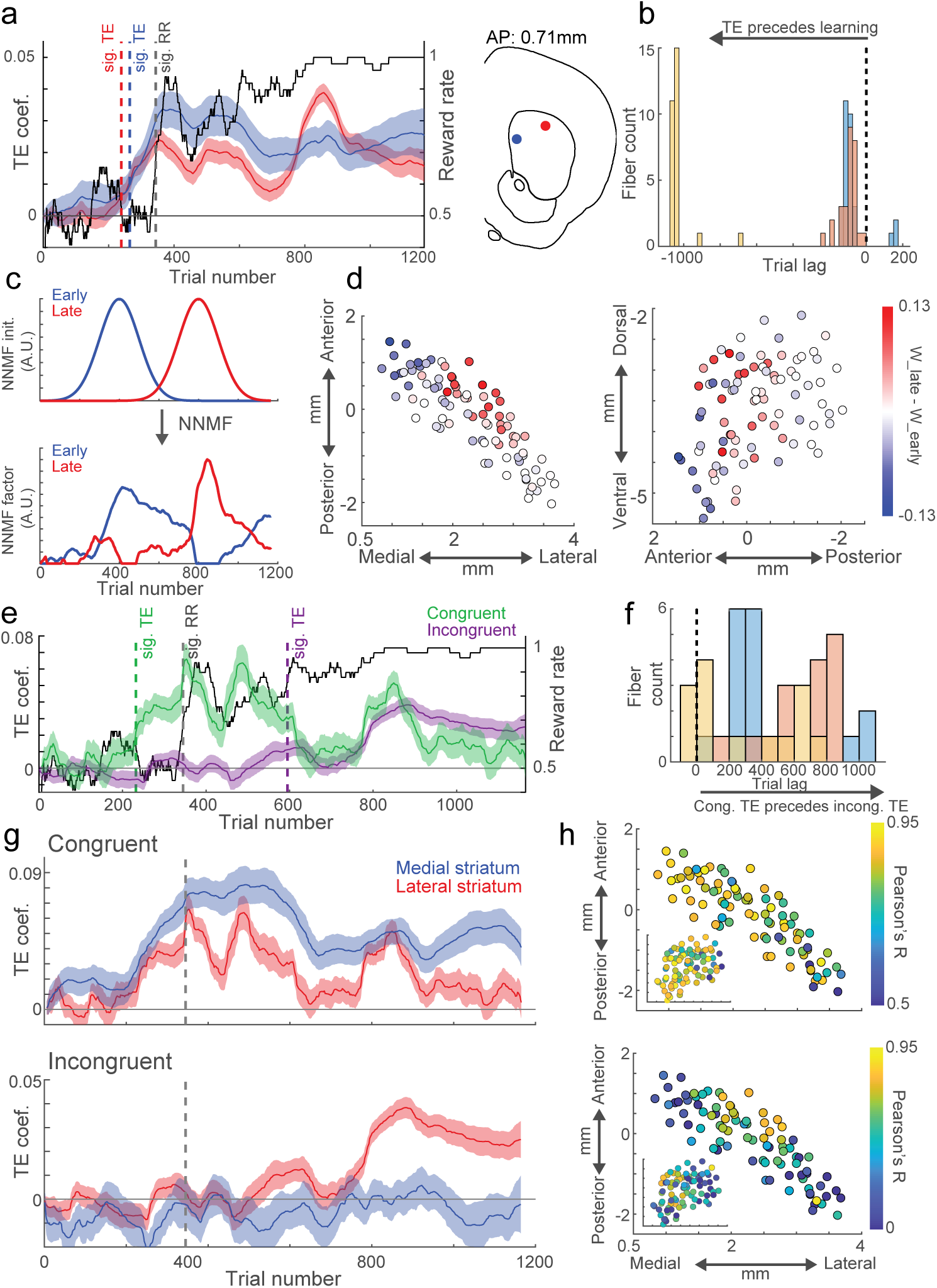
Emergence of TE encoding during learning precedes performance changes and follows distinct time courses for congruent and incongruent trials and across regions. **a,** Model estimates of trial by trial reward rates (black) and TE coefficients for two example fibers (blue, medial; red, lateral) across all trials and sessions through learning (see Methods). Dashed lines indicate the trials in which the TE coefficients and reward rates reached significance (p<0.01, shuffle test, see Methods). Fiber locations indicated in the coronal atlas schematic. **b,** Histogram of lags between the earliest trial where TE and reward rate reached statistical significance for each fiber in the three included mice. Colors indicate different mice. Negative values indicate that TE becomes significant prior to reward rate. **c,** Non-negative matrix factorization (NNMF) early and late component initialization (top) and output (bottom) for one example mouse (see Methods for details). **d,** Horizontal (left) and sagittal (right) plane maps of the differences in the weights for the late and early components of the NNMF for each fiber (circle). Positive values (red) indicate a stronger weight for the late component, and negative values (blue) a stronger weight for the early component (n = 88 fibers, 3 mice). **e,** Trajectory error coefficient estimates across congruent (green) and incongruent (purple) trials as in a for one representative fiber (the red site in a). **f,** Histogram as in b of lags between the earliest trial where congruent and incongruent TE reached statistical significance for each fiber. Positive lags indicate congruent preceding incongruent. Colors indicate different mice. **g,** Model estimates of trial by trial TE coefficients across trials as in a for congruent (top) and incongruent trials (bottom) for the two representative fibers in a (blue, medial; red, lateral). **h,** Correlations (Pearson’s) for each fiber of the TE coefficient estimates across all trials combined vs the TE coefficient estimates across all congruent (top) or incongruent (bottom) trials separately for each fiber (circle). Maps are shown in the horizontal plane, sagittal inset. Shaded regions in all plots are mean +/- S.E.M.

We tested whether the learning time course differed for positive (congruent) and negative (incongruent) TE signaling by fitting separate models for congruent and incongruent trials separately (Fig. 5e-h). At all locations, positive TEs emerged before negative (Fig. 5f), thus, rapid increases in positive TEs may drive the early component of the overall TE time course, while negative TEs may drive the late component. Indeed the overall TE time course was most strongly correlated with the positive TEs time course in the anterior medial striatum and with the negative TE time course in the central lateral striatum (Fig. 5g,h). Together, these results illustrate that dopamine trajectory error encoding evolves through different time courses across striatal regions which align with the rapid early emergence of positive TEs (anterior medial striatum) prior to learning and the later emergence of negative TEs (central lateral striatum) (see Discussion).

### Acetylcholine release encodes region-specific trajectory errors inversely to dopamine

Striatal acetylcholine is released primarily from tonically active interneurons^48^, which exhibit multiphasic dynamics to Pavlovian cues after learning^14,15^. Dopamine and acetylcholine release are proposed to exert contrasting effects on movement and learning, a hypothesis partly supported by their opposing signals *in-vivo*^16,49,50^. We asked whether cue-evoked acetylcholine release also encodes region-specific trajectory errors, and if so, how it relates to the properties of TE signaling in dopamine release. We used our optical fiber array approach to measure striatum-wide fluctuations in acetylcholine release with the genetically encoded sensor ACh3.0^51^ in 4 mice (Fig. 6a) in the loc + vis task (Fig. 1a). Acetylcholine release at cue onset at some locations displayed a tri-phasic profile consisting of a short latency peak, followed by a dip, then a later peak (or rebound), consistent with previous observations in rodents and primates^14,49,52–54^ (Fig. 6c). We quantified whether acetylcholine release scaled with TEs at each timepoint following the cue onset using an FIR model, as for dopamine (Fig. 3c, 6d), and found significant TE encoding across 33/112 (∼30%) locations (Fig. 6f). Interestingly, TE encoding was opposite in direction and peaked later in time than dopamine TE encoding (Fig. 6d-f). Acetylcholine TEs were sparser than for dopamine and were restricted primarily to locations in the anterior dorsal striatum (Fig. 6f). This region aligned well with the regions expressing the most prominent dips in dopamine release for incongruent trials (Extended Data Fig. 5a-c), which, together with the dopamine/acetylcholine timing relationships (Fig. 6e), suggest a potential mechanism for dopamine/acetylcholine interactions in the generation of negative TEs in acetylcholine release (see Discussion).

**Figure 6:**
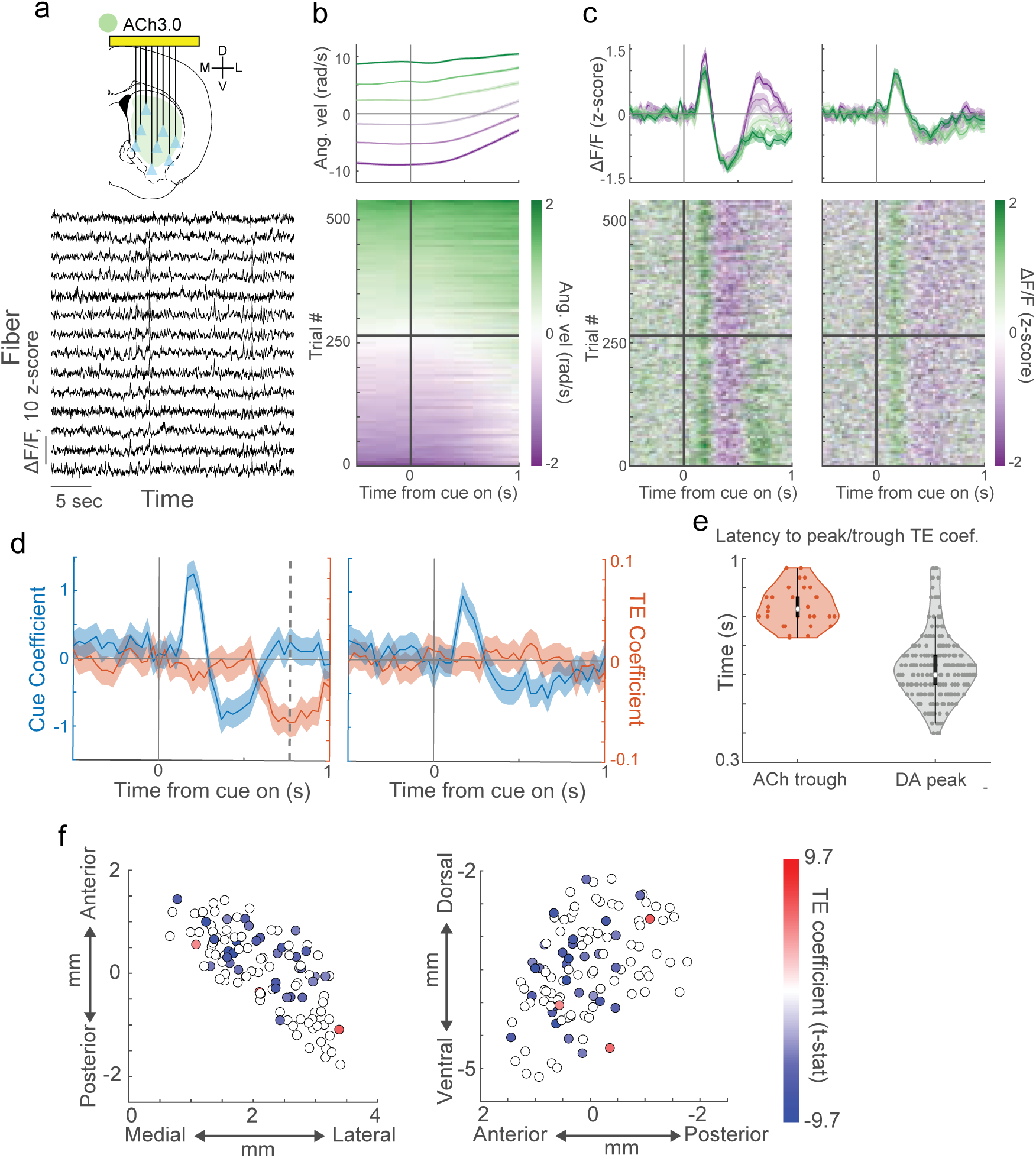
Trajectory error encoding in spatially organized acetylcholine release is inverse to dopamine. **a,** Top, schematic of striatum-wide acetylcholine release measurements with micro-fiber arrays. Bottom, example ΔF/F traces recorded simultaneously from 14 different fibers in a single mouse, ordered from dorsal (top) to ventral (bottom). **b,** Top, average congruence-signed angular velocity (+ congruent, - incongruent) aligned to cue onset for all congruent (green) and incongruent (purple) trials in one example mouse, split into thirds by magnitude (dark to light colors indicate fast to slow). Shaded regions are mean +/- S.E.M. Bottom, raster plot of signed angular velocity across all trials for the mouse at top, sorted by TE magnitude. **c,** Same as for b, but for ACh3.0 ΔF/F for two example fibers. Rasters are sorted by TE magnitude as in b, bottom. **d,** FIR model coefficients for cue (blue) and TE (orange). Shaded regions, 95% confidence interval. **e,** Violin plot across all fibers of the latencies to the minimum acetylcholine TE coefficient (orange), and maximum dopamine TE coefficient (gray). **f,** Horizontal (left) and sagittal (right) map of the most extreme TE (maximum, or minimum) coefficients for each fiber (circle). The color indicates the t-stat of the maximum coefficient. Open circles are not significant (n = 112, 4 mice; t-test on model coefficients, p<0.05 for 3 timepoints in a row within 1s cue window, Bonferroni corrected).

### Trajectory error signaling and choice can be modeled with reinforcement learning

Standard reinforcement learning models posit that dopamine release to cues encode reward prediction errors (RPEs) and emerge as a consequence of updates to a value function representing a temporally discounted expected future reward, driven by dopamine RPEs^8,13^. Values are calculated with respect to changing states, typically representing discrete cues and actions. We asked whether dopamine TE signals could be modeled under a temporal difference reinforcement learning (TDRL) framework, specifically as an actor-critic architecture (Extended Data Fig. 10)^13,55^. In our tasks, there was an inverse relationship overall between TE magnitude at cue onset and time to reward, consistent with temporal discounting (Extended Data Fig. 10a). However, TEs reflect conjunctive representations of cue identity and continuous movement, so discrete state representations of cues and actions separately would not contain the information necessary to reproduce dopamine TE signals. Therefore, we modeled a TDRL algorithm with a state representation composed of multiple neurons representing nonlinear functions of random weighted mixtures of cue identity and continuous velocity and position variables (Extended Data Fig. 10b). The state representation thus exhibited nonlinear mixed selectivity to task-relevant sensorimotor variables^56^. This representation was used to estimate state values and select actions (increases or decreases to angular velocity). Updates to the actor and critic were made based on a RPE, representing the dopamine signal. The random mixing of state variables supports value estimates and action probabilities that are based on combinations of cue and continuous movement variables necessary for TE encoding.

Our model reproduced accurate instrumental choice behavior with learning and also replicated key features of dopamine signals from our experimental results (Extended Data Fig 10c,d). Dopamine on all trial types initially rose to the cue, but signals were larger for congruent relative to incongruent trials and scaled continuously with TE magnitude. Surprisingly, the model also replicated the dopamine dip below baseline for negative TEs. This result cannot be explained merely as a consequence of temporal discounting. Rather, it arises from the interplay between the state representation and value updating in the RL algorithm. Specifically, since state representations at cue onset are correlated between congruent and incongruent states, positive value updates to congruent states also increase the value assigned to incongruent states. As a result, the critic systematically overestimates the value of incongruent states at cue onset. This overestimate is corrected when future state information (the agent’s action) further decorrelates the incongruent state from the congruent state, manifesting as a negative RPE, i.e. a dip in dopamine activity below baseline. These results suggest that a combinatorial encoding of sensorimotor variables in a continuous, high-dimensional neural state representation may allow TE signaling to arise naturally from the dynamics of RL.

## Discussion

We report anatomically organized encoding of trajectory errors (TEs) in striatal neuromodulator release which signal, when cues are present, whether, and how fast, animals are traveling in the correct or incorrect direction relative to a goal. TE encoding is relative to either mouse locomotion, cue visual flow, or both, enabling flexibility in situations when only certain reference frames are relevant (e.g. when direction of travel is signaled by visual flow rather than rotational locomotion). These signals are not driven by mouse acceleration or velocity per se and do not reflect predicted effort as they were observed when no locomotion adjustments were required (Fig. 3 and Extended Data Fig. 6). We propose that TEs act on striatal circuits to guide animals’ movement relative to landmarks in real time or across learning during goal-directed navigation, complementing cognitive-map based navigation in cortical and hippocampal circuits.

Error signals encoding mismatches between predicted sensory feedback and motor commands are well-documented across multiple species and brain regions and hypothesized to play roles in online error correction and learning^57–60^. Trajectory errors differ importantly in that they scale relative to arbitrary associations between sensory cues and movement directions and speed in visual and locomotor space, relative to goals. This property is essential to enable appropriate course corrections in landmark based navigation. Signals encoding negative errors in locomotion direction have been recently reported in a sparse subpopulation of posterior parietal cortex interneurons during navigation, but it is unclear how they could broadly impact learning or guide movement^61^. Trajectory error encoding in dopamine and acetylcholine release was widespread across the anterior striatum and thus positioned to powerfully impact striatal output and behavior across multiple parallel basal ganglia circuits^62^. Dopamine signals to learned cues or actions have been hypothesized to encode reward prediction errors (RPEs), which reflect the predicted magnitude, probability, spatial proximity, or timing of reward^21,22,24,29,37,63^. Unlike these signals, dopamine TE encoding incorporates the dynamic state of the animal or environment relative to a cued goal, information necessary to accurately learn and adjust behavior based on landmarks during goal pursuit.

Dopamine RPEs figure prominently in reinforcement learning models as teaching signals to modify representations of actions and cues associated with reward^8,9,13,64^. In temporal difference reinforcement learning (TDRL) models of Pavlovian and instrumental learning, dopamine RPEs to cues and actions emerge as a consequence of updates to a value function, representing expected future reward given the cues and actions (state). The TDRL framework has been widely invoked to explain the role of hippocampal and striatal networks in learning associations between cues (or locations), actions, and outcomes to guide navigation^1,65–67^. We found that continuous trajectory error signaling and mouse choice behavior could be reproduced in a TDRL model with a conjunctive state representation composed of neurons that represent nonlinear combinations of cue identity, velocity, and position tuning (Extended Data Fig. 10). Such a state representation could plausibly be generated by networks in frontal and motor cortical regions, which display mixed selectivity to locomotor and sensory variables and project to the striatum and midbrain dopamine neurons^56,68–70^. Individual midbrain dopamine neurons also can represent different combinations of sensory and movement parameters, suggesting that trajectory errors could be differentially computed across dopamine neuron subpopulations^38–40,71^. Once established, trajectory errors could modify striatal representations and thus impact future choice behavior. While these results show that TE signaling may be explained under a TDRL framework, alternative learning schemes are possible^72^. Rather than endorsing any single model, our results underscore the utility of expanding models and experimental paradigms to include conjunctive, continuous state spaces that capture the dynamics associated with naturalistic behaviors^64^.

TE signals could modulate the direction and speed of locomotion appropriately to reach goals, either on immediate (100s of ms) or long term (trials, days) timescales. Positive and negative TEs may respectively promote continuation or switching of ongoing behavior through bi-directional postsynaptic effects on striatal projection neuron (SPN) excitability and/or long term plasticity^73–75^. SPN firing can be modulated by artificial elevations of dopamine and acetylcholine signaling on 100s of ms timescales^76–78^, which, based on signal latencies relative to behavioral changes, may be fast enough to modulate the immediate probability of continuing or switching. However, rapid fluctuations in dopamine release may not strongly influence striatal output and behavior at physiological levels *in-vivo*^63,79,80^. Moreover, positive TEs consistently preceded changes in performance by many trials (Fig. 5 and Extended Data Fig 9), indicating that they alone do not directly drive immediate choice behavior but may participate in learning or indirect modulation of behavior in conjunction with other inputs. Negative TEs, in contrast, emerged after initial improvements in performance in some mice, indicating that they may participate in the later refinement of behavior. Interestingly, the relative magnitudes and changes during learning of positive and negative TEs were spatially organized, with late emerging negative TEs and dips below baseline most prominent in the anterior dorsolateral striatum (aDLS) and positive TEs more prominent in the anterior ventral and central striatum (Fig. 5, Extended Data Figs 2 and 5). These spatial differences may contribute to different forms of associative learning and behavior controlled by each region. For example, efficient task performance may involve parallel learning, by different subregions, of both the correct and incorrect cue-velocity associations to facilitate continuation and switching of actions, respectively. The late emergence of negative TEs in the aDLS may refine or crystallize behavior, consistent with a role of this region in habit formation^5,81^. Manipulations recapitulating the natural spatial and temporal properties of TE signaling^30^ will be needed to conclusively test their causal impact on striatal output signaling and behavior.

We found that trajectory error encoding in acetylcholine release was opposite in direction to dopamine and expressed across restricted territories, predominantly in the anterior dorsal striatum (Fig. 6). Striatal dopamine and acetylcholine have been proposed to oppositely impact movement^82^ and learning^17,18^ and display divergent dynamics to cues^14,16^. Cue-evoked dopamine release is often followed by dips in acetylcholine, and the magnitude and duration of the dips can be inversely modulated by dopamine through D2 receptors on cholinergic interneurons^49,50,83^. In our task, acetylcholine TE encoding lagged maximal dopamine TE encoding by several hundred milliseconds (Fig 6d,e), indicating that the dopamine TE signals may drive the inverse variations in acetylcholine. Increased dopamine release for positive TEs may suppress late components of cue-evoked acetylcholine, while decreased dopamine below baseline for negative TEs may disinhibit acetylcholine release by reducing persistent activation of D2 receptors. Interestingly, the acetylcholine TE encoding was more restricted to regions of the anterior dorsal striatum than dopamine, perhaps due in part to the different temporal dynamics of dopamine signaling in these regions. The more rapid signal decay kinetics in the anterior dorsal striatum, relative to ventral and posterior (Fig 1f-i) may facilitate the larger dopamine dips below baseline for incongruent trials (Extended Data Fig 2a-c) and stronger disinhibition of acetylcholine signaling. Region specific TE encoding in acetylcholine may perform a complementary role to dopamine in driving learning or choice by contributing to opposing changes in excitability or plasticity in different SPN subtypes^73,84^. This opposing influence may promote behavioral switching and the formation of negative cue-direction associations (i.e. cue → don’t go left) via specific striatal regions to drive efficient performance. Therefore, we propose that positive and negative dopamine trajectory errors promote region-specific positive and negative associations between sensory cues and dynamic behavior, which may support distinct aspects of landmark-based navigation across learning.

## Methods

### Animals

Male and female C57BL6 mice (18 total), ages 9-20 weeks at implantation were used across experiments. Mice were housed under a 12h light-dark cycle (light on 9pm-9am), and all experiments were performed during the dark cycle. Unimplanted mice were group-housed 2-5 per cage; after implantation, mice were singly housed. During behavioral training and imaging, animals were restricted to 1-2 mL of water per day so as to maintain 80-90% of initial body weight and otherwise had free access to food and water. All studies and procedures were approved by the Boston University Institutional Animal Care and Use Committee.

### Implant and surgeries

Multi-fiber arrays were fabricated in house according to the protocol in Vu et al., 2024. Briefly, 50 µm diameter fibers (46 µm core, 4 µm cladding, 0.66 NA) were threaded through holes in a custom 3D printed frame, measured to specific depths targeting the striatum, and glued in place using a UV-cured glue (Norland Optical Adhesive). Each implant had 23-114 fibers (Table S1), positioned in a pattern which targeted locations throughout the entire AP, ML, and DV extent of the striatum unilaterally (hemispheres counterbalanced across mice). The non-implanted ends of the fibers were bundled together and glued in a 0.8-1.3 diameter polyimide tube (MicroLumen), which was cut with a razor blade and polished using fine-grained sandpaper (5 µm followed by 3 µm, ThorLabs), such that the tops of each fiber were visible and smooth for imaging. Before implantation, a calibration procedure was performed to map the implanted fiber location with the imaging surface.

Pre-trained mice were outfitted with a custom metal head plate^31^ secured to the skull with Metabond (Parkell), several weeks before the implant and injection surgery. The headplate was lightly secured to enable removal. Following pre-training, mice were injected intracranially with AAVs carrying the genetically encoded fluorescent sensors dLight1.3b^33^ (pAAV-CAG-dLight1.3b, AAV9 or AAV5) or ACh3.0 (AAV9-hSyn-Ach3.0^51^) diluted 1:3 in PBS (final titer: 6×10^12, 4.25×10^12, or 3.75×10^12 GC/mL for dLight1.3b; 7.65×10^12 GC/mL for Ach3.0) and implanted with the prepared multi-fiber arrays. One mouse received the intracranial injections and implant during two separate surgeries. Mice were anesthetized under isoflurane (1-3%) and positioned in a stereotaxic frame (Kopf), then a large (∼4.5 x 3 mm) craniotomy was drilled over the left or right striatum (Table S1). AAV injections were targeted to 17-43 sites across striatum, chosen to maximize expression around the target fiber locations. Each site was pressure injected with 200-800 nL of the viral solution at a rate of ∼100 nL/min, using a pulled glass pipette. The array was then mounted on the stereotaxic frame, positioned relative to bregma using a reference fiber, and slowly lowered into the brain. The exposed brain was sealed with Kwik-Sil (World Precision Instruments), and a thin layer of tissue glue was applied to lightly seal the medial edge of the grid to the skull. C&B-Metabond (Parkell) was then used to secure the implant to the skull and seal any exposed bone and kwik-sil. A metal headplate, ring and plastic protective cap were secured with Metabond and a layer of Metabond blackened with carbon powder (Sigma) was applied to the exposed fibers and protective cap.

### Fiber localization

Fiber localization has been described in detail in Vu et al., 2024. Briefly, at the experiment end, mice were perfused transcardially with 1% phosphate buffered saline (PBS) followed by 4% paraformaldehyde. The implanted skulls were then prepared for CT scanning by exposing the ventral surface of the brain and soaking in a 25% Lugol’s solution contrast agent. The 3D CT-scans were registered to the Allen Mouse Brain Common Coordinate Framework 3D 10-micron reference atlas and the fiber tips were localized and mapped to the implant array design, using a combination of FIJI/ImageJ plugins and custom MATLAB apps developed in-house^30^. Atlas coordinate locations of all the fibers were generated via semi-automated fiber tip localization, and anatomical labels were assigned from the reference atlas. These labels were visually inspected and reassigned based on the boundaries in the CT image, when necessary for borderline cases. Only fibers localized to the striatum were included for analysis (Table S1). All fiber locations and spatial maps are plotted relative to bregma.

### Behavior

#### Head fixed behavior and visual cue display

To enable running in all directions, mice were headfixed over a hollow styrofoam ball treadmill floating in a 3D-printed plastic cradle with holes to deliver pressurized air^31^. Ball rotation was measured on 3 axes (pitch, yaw, and roll) with two optical mouse sensors (Logitech G203 mice with hard plastic shells removed, set to a 400 dpi sensitivity with a polling rate of 1 kHz), positioned at the ball equator at 90 degree angles from one another. Outputs from the optical mice conveying pitch and yaw rotation were converted to an analog voltage through a Raspberry Pi (3B+) at 100 Hz (maximum velocity of 3.5 m/s corresponding to output voltage of 3.3) and relayed into a NIDAQ board (National Instruments, PCIe6343). Velocity direction was read as a digital binary variable from the Raspberry Pi, through the NIDAQ board. For water reward delivery, a 60 mL syringe was connected with plastic tubing to a digitally gated solenoid valve (Neptune Research, #161T012), which was then connected to a post-mounted water spout. Licking was monitored by a capacitive touch circuit connected to the spout. Solenoid triggering and behavioral data acquisition was performed by the NIDAQ at 2kHz and synchronized and collected with custom MATLAB software.

Cues were designed using the Virtual Reality MATLAB Engine (ViRMEn^85^) and displayed on an array of 5 12×21 inch monitors arranged in a semi-circle approximately 13 inches in front of the mice. Each cue consisted of vertically or horizontally oriented black and white stripes extending 10 inches wide across the entire vertical monitor surface (21 inches). For four mice, the visual cues had two small white circles positioned symmetrically on both sides of the cue at an angle of pi/5.5 rad to indicate the trial end position. Angular (yaw) ball velocity in the loc + vis instrumental task (Figure 1) was converted into angular cue movement across the monitors in ViRMEn. Intertrial intervals with no cue present ranged from 7-14s, pseudorandomly drawn from a uniform distribution. Trials began with the onset of one of the two visual cues, presented at the center of the mice visual field. Each cue was presented pseudorandomly on 50% of trials, with no more than 3 trials in a row with the same cue. Cue rotation, trial end, and reward delivery were determined according to the task rules described below. After reward delivery or trial end, cues remained in place for 0.5 s before turning off. Task contingencies and trial structure were controlled via custom functions written in MATLAB.

After surgery, mice recovered for 2-6 weeks, allowing time for the fluorescent sensor to express. Three to five days before beginning behavioral training, mice were restricted to 1-2 mL water per day to maintain a body weight of 80-90% of their baseline body weight. For approximately one week prior to task training, mice underwent acclimation to the head-fixed setup for 10-30 minutes per day where unexpected water rewards (7 µL) were delivered at random time intervals drawn from a 5-30 s uniform distribution. Mice were then trained on a given task for 30-60 minutes per day, including 30 minutes of imaging time. After imaging sessions began, mice were trained and imaged with no more than 1-3 day breaks, except when switched to a new task. The task and training schedule for each task type is described below.

#### Loc + vis task

A total of 14 mice (8 M, 6 F; Table S1) for dopamine and acetylcholine measurements were trained on the loc + vis task (Figure 1a). Starting at cue onset, the cue moved laterally across the monitors with a speed and direction determined by the yaw velocity of the ball. Velocity (in rad/second) was converted into cue movement by a scaling factor of 0.02 to produce trial times with a median of 5 s (Extended Data Fig 10d). To match the correspondence between directional locomotion and cue movement relative to the mouse in natural scenarios, the direction of lateral cue movement was opposite to the running direction of the mouse (i.e. clockwise (right) ball rotations corresponded to positive velocities and clockwise (right) cue movement, which is experienced by the mouse as a counterclockwise (left) turn). The trial ended when the cue was rotated by pi/5.5 radians (∼33 degrees) in either direction. The cue pattern (horizontal or vertical stripes) indicated which trial end position (left or right) would be rewarded, and the mapping between cue and rewarded direction was counterbalanced across mice.

A subset of mice included in the loc + vis analysis were pre-trained on other task variants. No differences in TE signaling or spatial patterns were observed across mice with different pre-training schedules, so all were combined for analyses. Four of the dopamine mice were initially pre-trained on a ‘block task’ for 10-16 days with a similar structure to the loc + vis, but with only one visual cue, not presented in the loc + vis task (diagonal black and white stripes). Each running direction in the block task was rewarded at either 90-100% or 0-10% probability, and probabilities switched every 8-10 trials. Two of the dopamine mice were first trained on a ‘touch’ task in which the relationships between cue identity and direction were the same as the loc + vis task but choice direction and cue movement were determined by reaching movements to touch sensors. Four dLight1.3b mice were initially trained in the vis-only task (see below). The three mice included in the learning analysis (Fig 5 and Extended Data Fig 9) were pre-trained on the block task prior to the loc + vis. 3 ACh3.0 mice were pre trained on the loc + vis task before implantation, while one began training post-implant. Mice were imaged and trained on the loc + vis task until they reached a reward rate exceeding 80% within one session, followed by an additional 4-12 days. With the exception of the learning analysis (Fig 5), sessions where performance was lower than 80% were excluded from analyses.

#### Vis-only task

Four naive mice (2 M, 2 F) were introduced to the vis-only task (Figure 4a-h) prior to any training on other task variants. In this task, the relationships between visual cues and rewarded cue directions were the same as in the loc + vis task, but the cue movement was entirely independent of the animals’ running velocity. The cue direction and speed on each trial was yoked to trials from two mice who had previously completed the loc + vis task. Congruent and incongruent cue trajectories (speed + direction) on each trial were drawn randomly from a set of correct trials in the loc + vis task with durations less than 10s, which met the inclusion criteria (described in the data preprocessing section). The set of trials was chosen so that each cue was presented on 50% of trials and 70% of trials were rewarded and 30% unrewarded in each session. Reward was delivered when the cues reached the rewarded lateral position (same as loc + vis task) and omitted if the end cue position did not match the rewarded position. Imaging on the vis only task was carried out for 7-8 days after animals reached learning criterion (described in the data preprocessing section).

#### Loc-only task

After completing the loc + vis task, two mice (1M, 1F) were trained and imaged on the loc-only task (Fig. 4i-n). The task rules were the same as the loc + vis task, except that visual cue remained stationary at the center of the mouse’s visual field and did not move laterally with the ball velocity. The trial end and reward delivery were determined by calculating the theoretical position of the visual cue if it were coupled with the rotational velocity as in the loc + vis task. In other words, the locomotor requirements of the mouse were identical to the loc + vis task, but no visual flow was provided to indicate the mouse’s angular velocity and position relative to the goal. Mice were imaged on this task for 7-8 days with a performance above 80% rewarded trials.

#### Pavlovian task

Four mice were trained on a Pavlovian conditioning task in which a light cue was paired with a water reward (Extended Data Fig. 7). During each imaging session, mice received 60 presentations of a light and a tone cue (30 presentations each) in a pseudorandom order. Each cue was presented for a total of 6 seconds with a water reward delivered at a fixed delay of 3 seconds from cue onset, followed by an ITI randomly drawn from a uniform distribution of 4-40 s. Throughout each session, 8 random rewards were delivered during the ITI periods. Light cues were delivered with an LED (ThorLabs, M470L3, 470nm) with a 7 mW intensity, positioned ∼20 cm away from the mouse at a 45 degree angle contralateral to the implant. Tone cues (12 kHz, 80 dB) were delivered with a USB speaker positioned ∼30 cm away from the mouse. Mice were trained until a learning criterion, described in the data analysis section, was reached and then trained and imaged for an additional 2-10 days. Only trials in which the light was delivered were included in analysis.

**Table S1:** Mouse and Surgery Info.

| Mouse | Sex | Implant side | Fluorescent sensor | Fibers included/total | Task |  |  |  |
| --- | --- | --- | --- | --- | --- | --- | --- | --- |
|  |  |  |  |  | Vis-only | Loc + Vis | Loc-only | Pavlovian |
| 1 | M | R | dLight1.3b | 57/114 | ☐ | ☒ | ☐ | ☐ |
| 2 | M | R | dLight1.3b | 29/75 | ☐ | ☒ | ☐ | ☐ |
| 3 | M | L | dLight1.3b | 28/52 | ☐ | ☒ | ☐ | ☐ |
| 4 | M | R | dLight1.3b | 31/51 | ☐ | ☒ | ☐ | ☐ |
| 5 | F | R | dLight1.3b | 40/71 | ☐ | ☒ | ☒ | ☐ |
| 6 | M | L | dLight1.3b | 45/70 | ☐ | ☒ | ☒ | ☐ |
| 7 | F | R | dLight1.3b | 24/50 | ☒ | ☒ | ☐ | ☐ |
| 8 | M | L | dLight1.3b | 40/53 | ☒ | ☒ | ☐ | ☐ |
| 9 | F | R | dLight1.3b | 32/51 | ☒ | ☒ | ☐ | ☐ |
| 10 | M | L | dLight1.3b | 30/51 | ☒ | ☒ | ☐ | ☐ |
| 11 | F | R | Ach3.0 | 8/23 | ☐ | ☒ | ☐ | ☐ |
| 12 | F | R | Ach3.0 | 47/67 | ☐ | ☒ | ☐ | ☐ |
| 13 | F | R | Acg3.0 | 18/23 | ☐ | ☒ | ☐ | ☐ |
| 14 | M | R | Ach3.0 | 43/97 | ☐ | ☒ | ☐ | ☐ |
| 15 | F | R | dLight1.3b | 18/103 | ☐ | ☐ | ☐ | ☒ |
| 16 | F | R | dLight1.3b | 53/84 | ☐ | ☐ | ☐ | ☒ |
| 17 | F | R | dLight1.3b | 51/74 | ☐ | ☐ | ☐ | ☒ |
| 18 | M | R | dLight1.3b | 27/74 | ☐ | ☐ | ☐ | ☒ |

### Optical measurements

Fluorescence around the implanted tips of each fiber was measured by imaging the polished surface of the fiber bundle with a table mounted microscope (see Vu et al. Neuron, 2024 for additional details). Excitation light was provided by high power LEDs (470nm and 405nm; Thor labs, No. SOLIS-470C, SOLIS-405C). Light power from the objective was adjusted to be within the range of 60-75 mW, resulting in a power of 1.4∼1.7mW/mm2 at the fiber tips. Emission light was bandpass filtered (Chroma, No 525/50m) and focused with a tube lens (Thor labs, No TTL165-A) onto the CMOS sensor of the camera (Hamamatsu, Orca Fusion BT Gen III). Imaging data was acquired using HCImage live (HCImage live, Hamamatsu). Single wavelength 470nm excitation was carried out with continuous imaging at 30Hz (33.33ms exposure time). In 7 mice, quasi-simultaneous excitation with 405nm and 470nm light was conducted for 1-2 days to address potential artifactual contributions (Fig. 2b, Extended Data Fig. 3). While increases in dopamine binding still results in slight (negative) fluorescence changes with 405nm illumination, no significant TE encoding was present (Extended Data Fig. 3, 0/236 fibers significant for 405, 103/246 significant for 470, t-test on model coefficients, p<0.05 for 3 timepoints in a row within 1s cue window, Bonferroni corrected for number of timepoints). Excitation wavelength separation was achieved by alternating the 405 and 470nm LEDs and synchronizing each with imaging acquisition. Alternation was performed at 36 Hz to achieve a frame rate for each excitation wavelength of 18 Hz (20ms exposure time).

### Analysis

Shuffle tests for comparison of two groups (Fig. 2c, 3h, 4h,n, 5a, Extended Data Fig. 3d, 5b, 6c,j) were performed by shuffling group identities and taking the difference of the means of the two groups 10,000 times to obtain a null distribution. For one-sided tests, significance was determined if the true difference of the means exceeded the 99th percentile of the null distribution. For two-sided tests, significance was determined if the true difference was outside of the 0.5-99.5 percentile of the null distribution.

#### Behavioral variables and session inclusion criteria

Angular velocity was calculated from ball yaw velocity, converted into rad/s, and signed according to one of three conventions, as indicated in each plot: 1) according to direction (+ left, - right), 2) according to direction relative to the implant (implant-signed, + contra, -ipsi), and 3) according to congruence (congruence-signed, + congruent, - incongruent). Linear velocity was defined based on the pitch and roll ball velocities, converted into m/s and combined so that

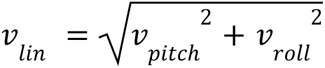

Linear and angular velocities were smoothed using a lowess filter with a smoothing window of 0.3 s, then low pass filtered with a 1.5 Hz cutoff. Linear acceleration was the derivative of linear velocity. Ipsilateral and contralateral acceleration was computed as the derivative of absolute value of the angular velocity, separated into periods where the mouse velocity direction was either ipsilateral or contralateral to the implant (Extended Data Fig. 6e,f). Angular acceleration was computed as the derivative of the implant-signed angular velocity (+ contralateral, - ipsilateral), such that positive values indicate increasing velocity in the contralateral direction or decreasing velocity in the ipsilateral direction, and negative values indicate decreasing velocity in the contralateral direction or increasing velocity in the ipsilateral direction. Linear and angular acceleration were smoothed with a lowess filter with a smoothing window of 0.3 s, and lowpass filtered with a threshold of 1.5 Hz.

Congruence was defined based on whether the yaw velocity direction was towards (congruent) or away from (incongruent) the rewarded (goal) position. Trajectory error was defined as the magnitude of the angular velocity (speed) in rad/s, signed by congruence (congruent, +; incongruent, -). The trajectory error and congruence on each trial of the loc+vis and loc-only tasks was defined as the average angular ball velocity within a 0.3 second window before cue onset. This window was chosen because it was before the average velocity began to change in response to the cue. For the vis-only task, the cue based trajectory error and congruence on each trial were computed based on the average angular velocity of the cue within a 0.2 second window after cue onset.

Except for the learning analysis (Figure 5 and Extended Data Figure 9), only days in the loc + vis and loc-only tasks where performance was above 80% correct were included (3-10 days). To ensure that we only included trials where the mice were following the task rules, incorrect trials and trials where there were more than 2 incorrect trials within a window of 10 surrounding trials were excluded. Finally, trials were excluded in which the total velocity, computed by taking the square root of the sum of squares of the ball velocity (in m/s) in all three dimensions (pitch, yaw, roll) and smoothed as for linear velocity described above, dropped below 0.3 m/s within a half second before cue onset, indicating that the mouse was at rest. This threshold was selected manually based on the distribution of total pre-cue velocities. For the vis-only task, a lick index was calculated to assess learning of the cue-direction associations (Figure 4a-c) by comparing the sum of the licking 1 second before reward (*L*_*rew*_) and 1 second before cue onset 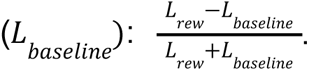 Days were included which fit two criteria: 1) the average lick index for rewarded trials was significantly greater than zero, and 2) the average lick index for rewarded trials was significantly greater than unrewarded trials (6-7 days, p<0.01, one-sided shuffle test with 10,000 iterations). For criteria 1, the average lick index was compared to an empirical null distribution, computed by randomizing the label for reward and baseline time periods. For criteria 2, the difference in lick index between rewarded and unrewarded trials was compared to an empirical null distribution by randomizing the trial type label and computing a new lick difference 10,000 times. For the pavlovian task, the lick index was computed as 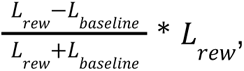 and normalized to the maximum single-trial lick index across mice. Days in which this lick index was below 0.15 were excluded.

#### Signal pre-processing, ΔF/F extraction, and fiber inclusion criteria

Movies of the fiber bundle surface were processed to extract changes in fluorescence from each fiber as described in detail in Vu et al., 2024. Movies were first motion-corrected using a cross correlation algorithm described previously^30^. Mean fluorescence time series were extracted from circular regions of interest (ROIs, ∼25 µm diameter) drawn within each fiber. Relative change in fluorescence ΔF/F was calculated from the mean fluorescence time series by subtracting the 8th percentile fluorescence over a 30s sliding window.

Fibers were included in analysis if they were conclusively determined to be in the striatum (see Fiber Localization above) and if they exhibited a significant positive or negative ΔF/F at cue onset. Significance was determined by whether the baseline corrected cue triggered average ΔF/F within the first second after cue onset was higher or lower than the 99% confidence interval of a null distribution for at least 3 timepoints in a row. The null distribution was determined for all analyses by calculating baseline corrected triggered averages from randomly selected timepoints in the task over 10,000 iterations. Baseline correction was performed on each trial by subtracting the average ΔF/F within a 0.5 second window before cue onset. Baseline correction was performed for all analyses included in the paper to ensure that we were reporting relative changes at cue onset, but the main results were similar with and without baseline correction (Extended Data Fig. 4).

#### Temporal dynamics

To assess the timescales of dopamine signaling (Fig 1f-i), we constructed autocorrelograms for each recording location by computing pearson’s correlation coefficients, using the corrcoef function in MATLAB, on the full ΔF/F trace concatenated across days with >75% correct performance with time shifts up to 2 s. Resulting autocorrelograms were normalized to the maximum correlation coefficients. To determine the decay constant (Fig. 1i), we fit each autocorrelogram to an exponential model using the ‘fit’ function in MATLAB:

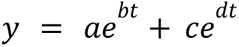

Where *y* is the autocorrelogram trace, and *t* is the time shift in seconds. The decay constant was determined as the minimum of *b* and *d*, indicating the faster decay term. The second, slower decay term accounts for noise, expressed as positive correlations at longer time lags. Autocorrelograms in Fig 1h have the noise term subtracted out.

#### Congruence and trajectory error models

To compute congruence encoding at each recording site (Fig. 2), ΔF/F traces, concatenated across included sessions, were modeled as a linear combination of finite impulse response functions (FIRs) at each timepoint (based on imaging sampling rate) within a window around each cue onset, representing trial level variables, including congruence, cue identity, linear acceleration and directional angular acceleration (Figs 2d,e and 3c). 4 mice with a strong directional bias which only ran in one direction at cue onset were excluded to avoid covariance between congruence and cue identity. The following function was fit to estimate a coefficient at each time point around the cue onset using matlab’s fitlm function:

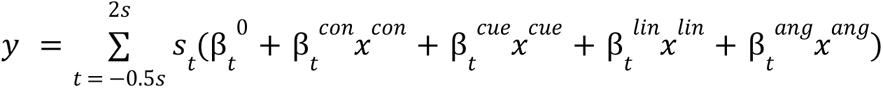

Where *s*_*t*_ is 1 at time t from cue onset and zeros everywhere else, to create a cue kernel for each variable. *x^con^* is 1 on congruent trials and -1 on incongruent trials, defined above. *x^cue^* is 1 on contralateral cue trials and -1 on ipsilateral cue trials to allow for possible different cue responses depending on cue identity. *x^acc^* is the minimum linear acceleration within the first 0.4 s of cue onset to account for rapid, transient decelerations after cue onset. *x^ang^* is the average angular acceleration from 0.5 to 1 s after cue. Including the acceleration terms as trial-level FIR functions allows for potential temporal delays in the relationship between dopamine release and movement with the assumption that the largest acceleration within a given window will have the strongest effect on the signal. Trajectory error (TE) encoding for each recording site (Fig. 3) was calculated by modeling ΔF/F as in the congruence model, except the congruence terms (β*_t_^con^ x_t_^con^*) were replaced by trial-by-trial scaling TE terms (β*_t_^TE^ x_t_^TE^*) defined by the average angular velocity within a 0.3 second window before cue onset, with the sign indicating whether the velocity direction is congruent or incongruent. To compare different trial types (congruent vs incongruent, or contra vs ipsi), we included two separate sets of trajectory error FIRs which represented the TE on each trial type. For comparison models, four mice with a strong directional bias which only ran in one direction at cue onset were excluded to avoid covariance between congruence and cue identity. For the vis-only task (Fig. 4), two sets of TE FIR terms were included, one based on visual cue movement (cue TE) and one computed based on angular velocity (loc TE) (Fig. 4g). For the loc TE FIR, TE was defined as the magnitude and direction of the animal’s angular velocity within 0.3 s before cue onset, relative to the rewarded direction of the cue movement, if the cue movement were determined by angular velocity as in the loc + vis task. The cue TE was defined as the average direction and magnitude of visual cue velocity on each trial relative to the rewarded direction, within a 0.2s window after cue onset. To compare between the loc-only and loc + vis task, data from the loc + vis task were fitted with a model in which the TE terms were replaced with cue TE terms, computed the same way as the cue TE in the vis only task. For the pavlovian task (Extended Data Fig. 7), the TE terms were replaced by trial-by-trial angular velocity magnitude terms (average angular velocity magnitude within a 0.3 s window before cue onset).

Significant model coefficients (Fig. 2g, 3e, 3d, 6f; Extended Data Fig. 5, 7c, 8c-d) were determined using a t-test on the FIR coefficients within a 1 s cue window. Fibers were significant if p<0.05 for 3 timepoints in a row, Bonferroni corrected for the number of timepoints within the 1 s window.

#### Spatial gradient analysis

To determine the anatomical axes of maximal spatial variation in model coefficients and the gradients along each dimension in congruence and trajectory error encoding (Figs. 2 and 3), we fit a model to describe the maximum of the FIR coefficient t-statistic after cue onset as a function of the fiber location in each dimension, relative to bregma. This can be written as:

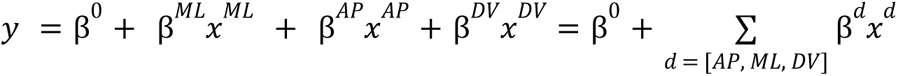

Where *y* is a 1 x n vector of the maximum coefficient t-statistics for each of n fibers. *x* is the position of each fiber in the ML, AP, or DV axis. The axis of maximum variance is defined by [β*^ML^*, β*^AP^*, β*^DV^*]. To compute differences in the spatial gradients between the encoding of two different variables, we modeled the two sets of coefficients across fibers as a function of their location in each dimension, with an added interaction term between coefficient type and recording location. Significance in each dimension, *d*, was determined based on the p value of the interaction coefficient β*^ds^*:

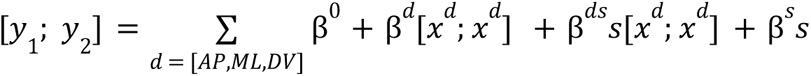

Where *y*_1_ *and y*_2_ are the two sets of coefficients. Here, *s* is a categorical variable with 0s for the data points corresponding to *y*_1_ and 1s for data points corresponding to *y*_2_.

#### Acceleration correlations in the intertrial interval

To determine the relationship between dopamine release and acceleration independent from the task, we modeled ΔF/F traces during the ITI periods starting from one second after reward delivery to the next cue onset as a function of linear and angular acceleration at different time shifts. The ΔF/F trace was first high pass filtered with a 0.8 Hz threshold to remove slow fluctuations that could be the result of a hemodynamic artifact. A separate model was fit for frame-by-frame temporal offsets between ΔF/F and ipsilateral and contralateral acceleration from -0.5 to 0.5 seconds. At each time lag the following model was fit:

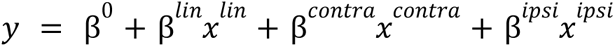

Where y is the the highpass filtered ΔF/F, *x^lin^* is linear acceleration, *x^contra^* is the contralateral acceleration during time periods when the angular velocity is in the direction contralateral to the implant, 0 otherwise, and *x^ipsi^* is ipsilateral acceleration during time periods when the angular velocity is in the direction ipsilateral to the implant, 0 otherwise. *x^ipsi^* and *x^contra^* are positive when the angular velocity is increasing and negative when the angular velocity is decreasing in either direction. This is in contrast to the implant-signed angular velocity described previously.

#### Learning analysis

To determine how dopamine TEs evolve across learning, we performed a dynamic regression analysis adapted from the PsyTrack approach^47^. Briefly, the magnitude of the dopamine cue response during a 0.2 s window around the time of maximal TE encoding (fig 3h) was modeled as linear-Gaussian model of the trajectory error *x_t_*,

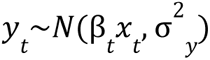

where the TE coefficients, β*_t_*, change on each trial based on a Gaussian random walk,

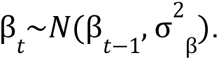

The smoothness hyperparameter, σ_β_, governs the rate of change of the TE coefficients as detailed in Roy et al 2021. We fit this model separately to data from each fiber. The time course of β*_t_* was inferred from the length *R* sequence of dopamine magnitudes and TEs, where *R* is the number of trials across learning. β*_t_*, σ^2^*_y_*, σ^2^_β_ was learned for each fiber using *maximum a posteriori* (MAP) estimation. The smoothness hyperparameter was determined by grid search: the MAP estimate of β*_t_*, σ^2^*_y_*, was computed for ten values of σ^2^_β_ in a fixed range (10^-4^ and 10^-1^) and the model with the highest test log-likelihood was selected. Five-fold cross-validation was performed to find the optimal model for each fiber. The MAP estimate was determined using a quasi-Newton method (BFGS) from the python *scipy* library. Further details of the PsyTrack dynamic regression approach can be found in Roy et. al 2021.

The dynamic TE coefficient was considered significant when the lower bound of the 99% confidence interval surpasses 0. Performance was considered significant when the reward rate, computed with a 50 trial sliding window exceeds the 99% confidence interval of an empirical null distribution computed by shuffling the trial number labels and recomputing the sliding window reward rate 10000 times.

#### TD RPE model simulation

A reinforcement learning agent was trained on a stimulus response task similar to the task from Figure 1. At each time step, the agent selects one of two actions: accelerate left or accelerate right. Based on the selected action, the angular velocity (ω) at each time step is determined by the following equations:

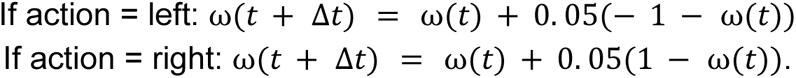

Thus, whether and how much the velocity changes is determined by the current velocity, and the direction of the change is determined by the selected action. Here *v* < 0 indicates right velocities and *v* > 0 indicates left velocities. These equations bound the velocity by +1 and -1, and the maximum acceleration at 0. 05/Δ*t*.

The state input used for action selection and learning (*s*_0_) includes 10 units each representing evenly tiled Gaussian tuning curves, for both the continuous position and velocity variables (separate units for position and velocity). 3 units represent the binary cue identity, or goal, as a categorical variable: one unit for the left cue, one unit for the right cue, and one unit for no cue. These 23 variables are provided as inputs with random weights to a second layer of N = 1000 neurons, according to the following equations:

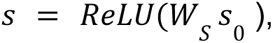

where *W*_*s*_ is a N×23 encoder matrix drawn randomly from a standard normal distribution, and ReLU is the rectified linear function, which maps negative inputs to zero and otherwise leaves them unchanged.

At each timestep, the action is selected according to probabilities given by a softmax function of the actor output:

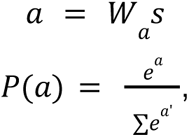

Where *W_a_* is a 2×N matrix representing the “actor” weights, updated at each timestep with a learning rate of α = 0.25/N using the following equation:

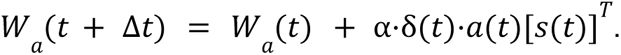

δ(*t*) represents the temporal difference reward prediction (tdRPE), which models dopamine release and is computed as follows:

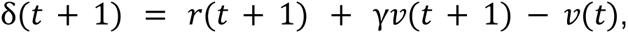

where *r* indicates the presence (1) or absence (0) of reward, γ = 0. 9 sets the temporal discount factor, and *v*(*t*) is the value function determined by the following equations:

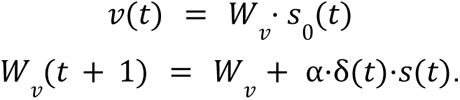

*W*_*v*_ is a vector of “critic” weights used to estimate value. On each trial, the position (*x*) is initialized at *x*_0_ = 0, and the velocity (ω) is initialized randomly based on a normal distribution. The position is updated on each timestep based on the velocity, according to *x*(*t* + 1) = *x*(*t*) + ω(*t*). One of two visual cues (left or right) is presented randomly with 50% probability at timestep 5, and the trial finishes when the agent reaches the end position *x* =± 1. If the left cue is present, the agent will be rewarded for reaching *x* =+ 1, for the right cue, the agent will be rewarded for reaching *x* =− 1.

## Acknowledgements

This work was supported by a Klingenstein-Simons’s Foundation fellowship, Whitehall Foundation Fellowship, National Institute of Mental Health (R01 MH125835) to M.W.H; NIMH F31NS127536-01A1 to E.B; NIMH F32MH120894 to M.-A.T.V; JSTPN Early Stage Training in Neuroscience Award to Y.Z.. We thank the Boston University Centers for Neurophotonics and Systems Neuroscience for financial and technical support, Micro CT core, especially Sydney Holder, for providing equipment and technical expertise for micro-CT scanning and Boston University Animal Science Center for providing central laboratory and animal care and support resources. We thank Drs. Lin Tian and Yulong Li for providing dLight1.3b and Ach3.0 viral vectors and Drs. Chantal Stern and Michael Hasselmo for feedback on a draft of the manuscript.

---

**Extended Data Figure 1:**
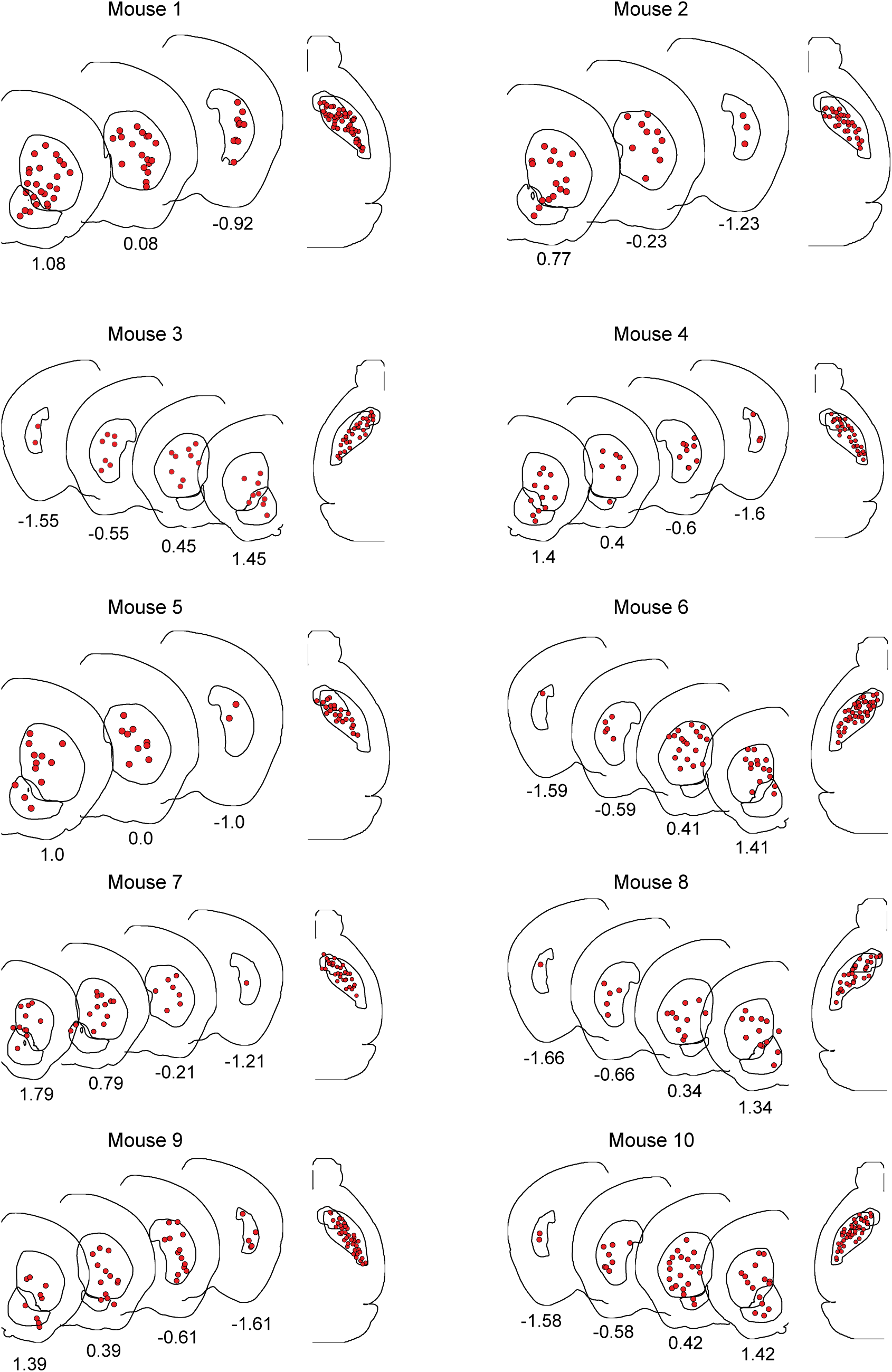
Locations of all fibers across individual mice. Atlas registered locations of all fibers (red circles) included in analysis for each of 10 mice plotted in the coronal and horizontal planes relative to atlas common coordinate framework (see Methods). Each coronal plane represents a slice with a 1 mm thickness. Numbers beneath each coronal section are AP coordinates of the most anterior section of the slice, in mm relative to bregma. A few fibers appear outside of the striatal boundaries (i.e. in mice 7, 8 and 9), due to slight deviations in the actual striatum boundaries of individual mice, as determined by the CT image, from the atlas striatum boundaries.

**Extended Data Figure 2:**
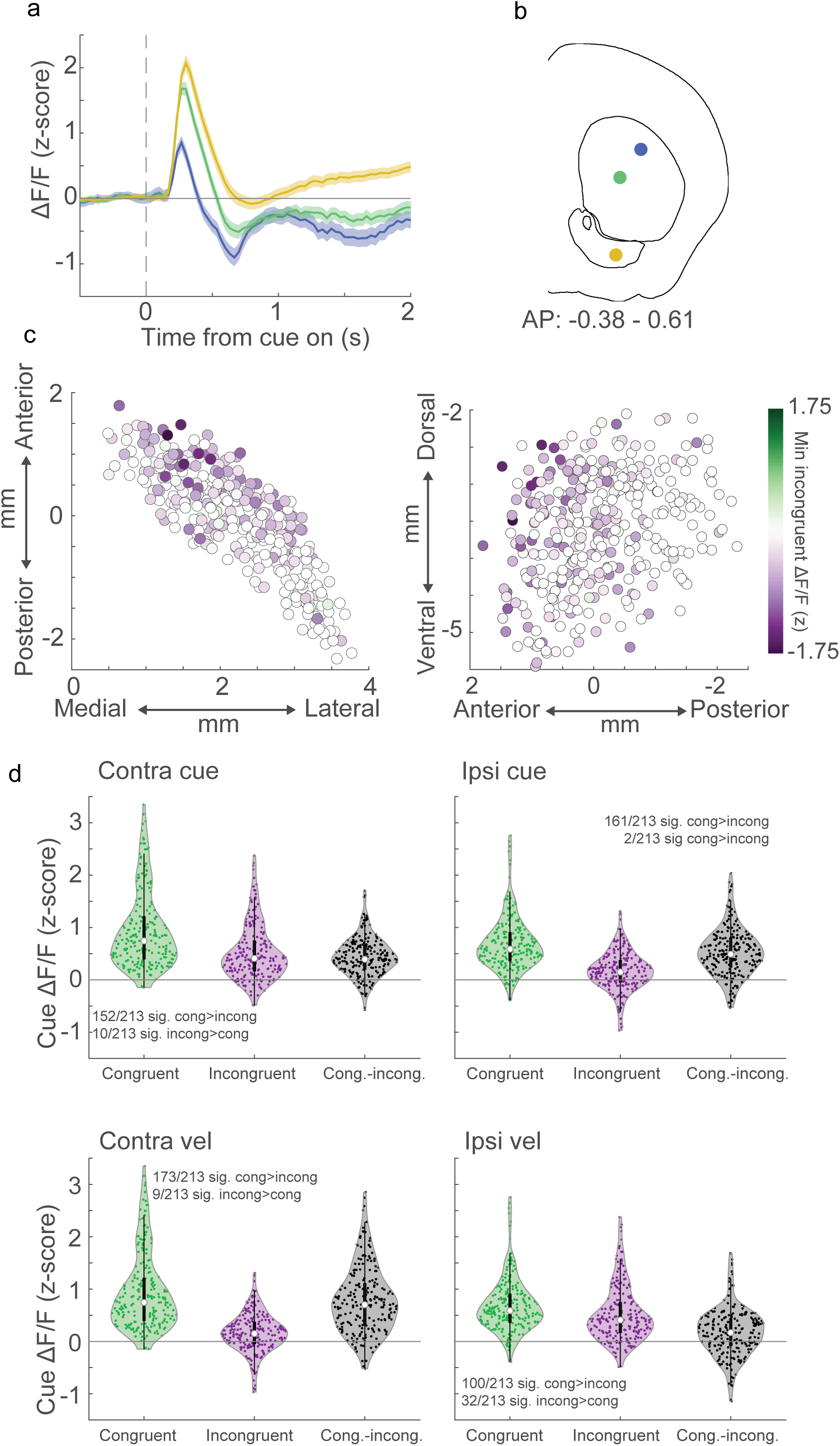
Spatial organization of dopamine dips on incongruent trials and consistency of congruence encoding across trial types. **a,** Average baseline-corrected ΔF/F aligned to cue onset on all incongruent trials for three simultaneously recorded example fibers in one mouse. Shaded regions are 95% confidence intervals. **b,** Coronal section showing the fiber locations for the example traces shown in a. **c,** Horizontal (left) and sagittal (right) maps of the minimum of the mean z-scored ΔF/F across all incongruent trials for each fiber (circle) across all mice (n = 351 fibers, 10 mice). Empty circles are fibers with no significant dip below baseline (mean < 99% confidence interval of empirical null distribution for 3 consecutive timepoints). **d,** Violin plots of average cue responses from 0.5-1s after cue onset across fibers and mice for congruent trials (green), incongruent trials (purple), and the difference between congruent and incongruent trials (grey), for contralateral (left), and ipsilateral (right) cue presentations (top) and pre-cue angular velocity directions (bottom) (n = 213 fibers, 6 mice).

**Extended Data Figure 3:**
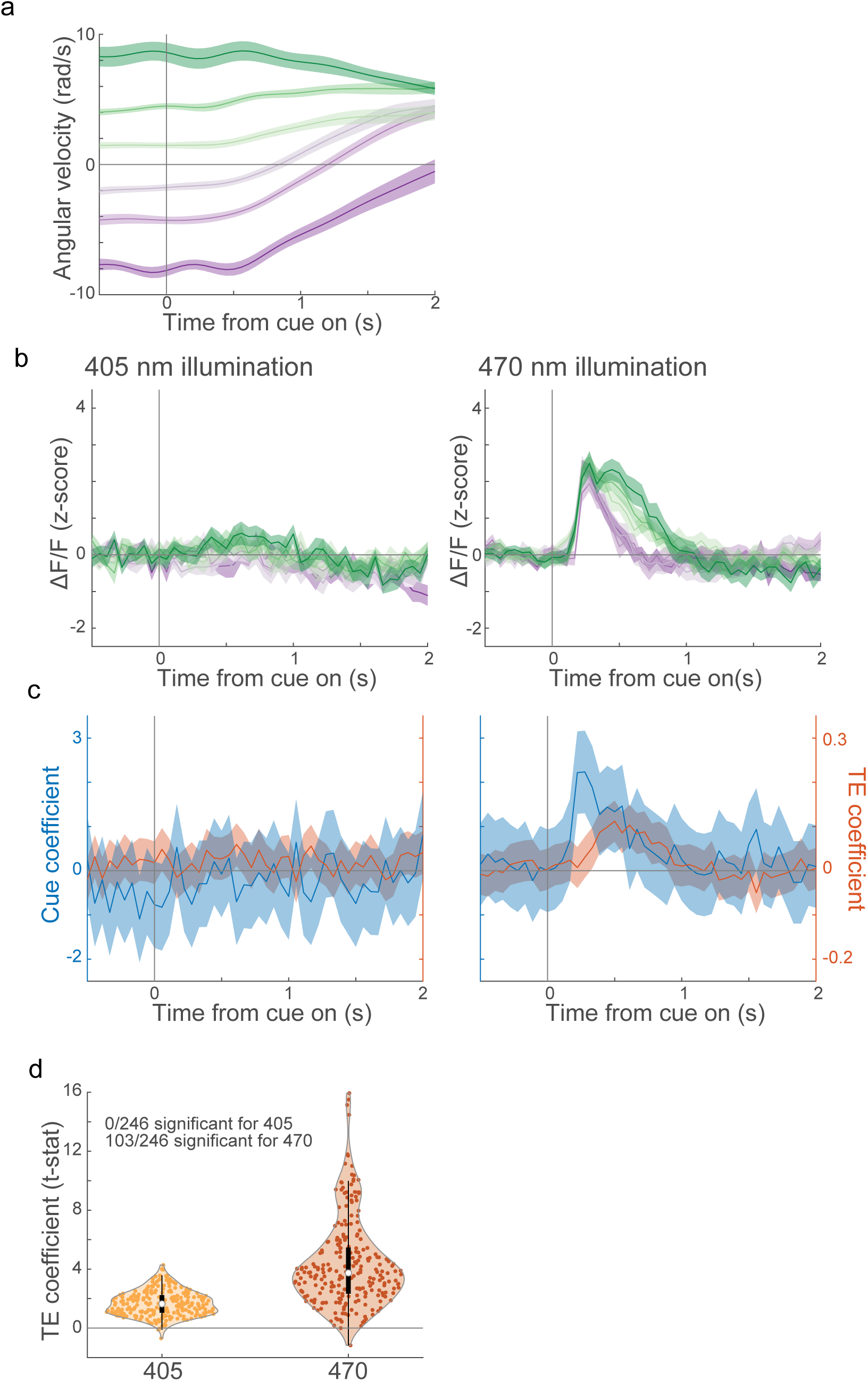
Trajectory errors are not a result of movement or hemodynamic artifact. **a,** Average congruence-signed angular velocity (+ congruent, - incongruent) aligned to cue onset for congruent (green) and incongruent (purple) trials, split into thirds by velocity magnitude (dark to light colors indicate fast to slow) for one recording session with pseudo-simultaneous recordings with 405 nm and 470 nm excitation. Shaded regions, mean +/- S.E.M. **b,** Same as a, but for ΔF/F from the 405nm (isosbestic) (left) and 470 nm (right) excitation. **c,** Cue (blue) and trajectory error (orange) coefficients for the 405 nm illumination isosbestic control (left) and 470 nm illumination (right). Shaded regions, 95% confidence intervals. **d,** Violin plot showing the distribution of the maximum TE coefficient across fibers and mice (n = 246 fibers, 7 mice, significance for each fiber determined by t-test on model coefficients, p<0.05 for 3 timepoints in a row within 1s cue window, Bonferroni corrected).

**Extended Data Figure 4:**
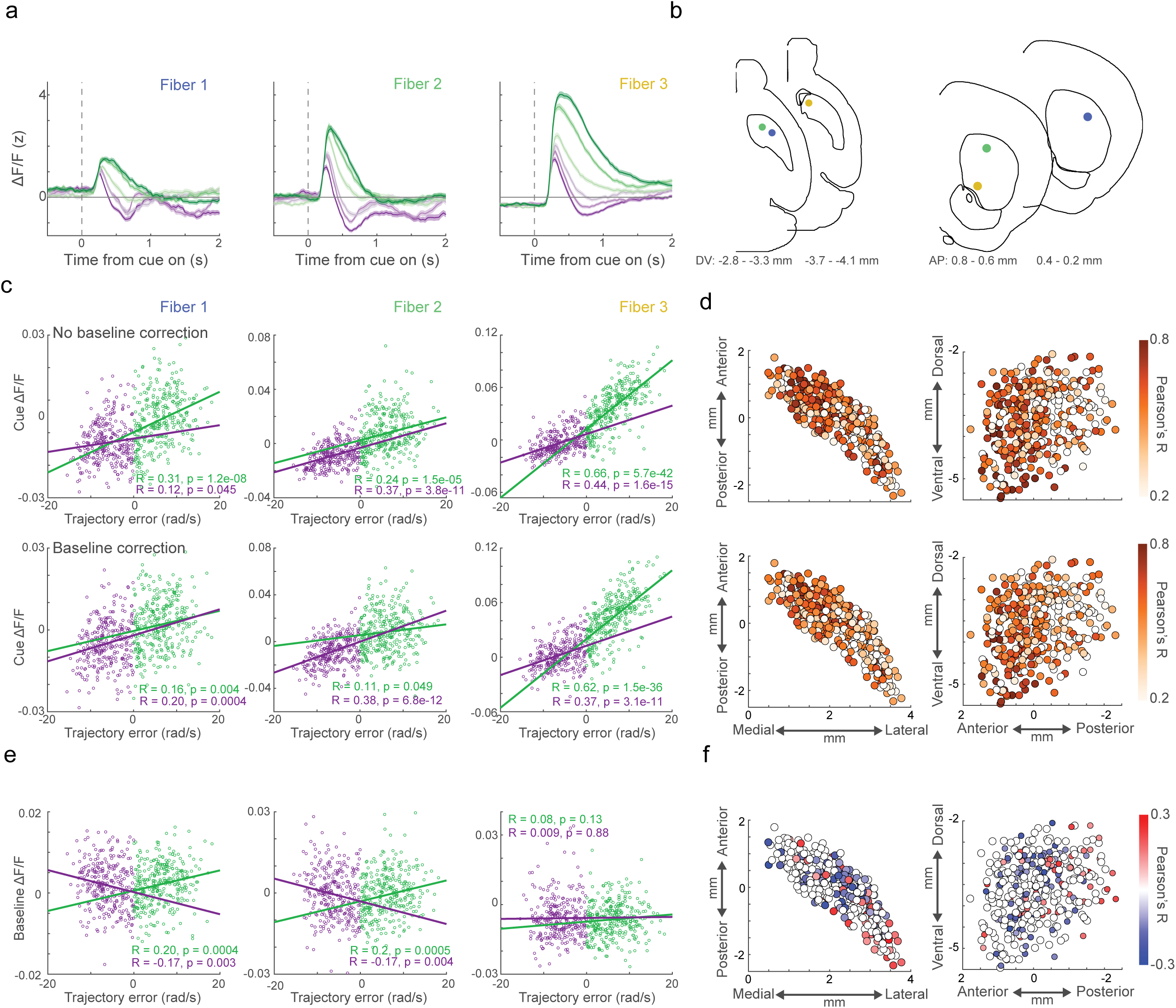
Trajectory error encoding is continuous across angular velocities and is not affected by differences in pre-cue baseline signals. **a,** Average ΔF/F (no baseline correction), for three example fibers from one mouse, aligned to cue onset for congruent (green) and incongruent (purple) trials split into thirds by magnitude (dark to light colors indicates fast to slow). Shaded regions, mean +/- S.E.M. **b,** Horizontal (left) and coronal (right) atlas sections showing the locations of the examples in a. **c,** Scatter plots of average ΔF/F vs TE within the 0.5 to 1 second window after cue onset on congruent (green) and incongruent (purple) trials with (bottom) and without (top) baseline correction. Each circle is a single trial for the fibers shown in a. Lines show the best linear fit from least-squares regression for congruent and incongruent trials separately. Inset text: Pearson’s correlation coefficients and p-values. **d,** Horizontal (left) and sagittal (right) maps of Pearson’s correlation coefficients between trajectory error and average cue ΔF/F across all trials, with (bottom) and without (top) baseline correction for each fiber (circle) across mice. Open circles are not statistically significant (p<0.01, n = 351 fibers, 10 mice). **e,** Same as c but for the baseline ΔF/F (average within 0.5 seconds before cue onset). **f,** Same as d but for correlations between absolute value of trajectory error (angular velocity magnitude) and baseline ΔF/F.

**Extended Data Figure 5:**
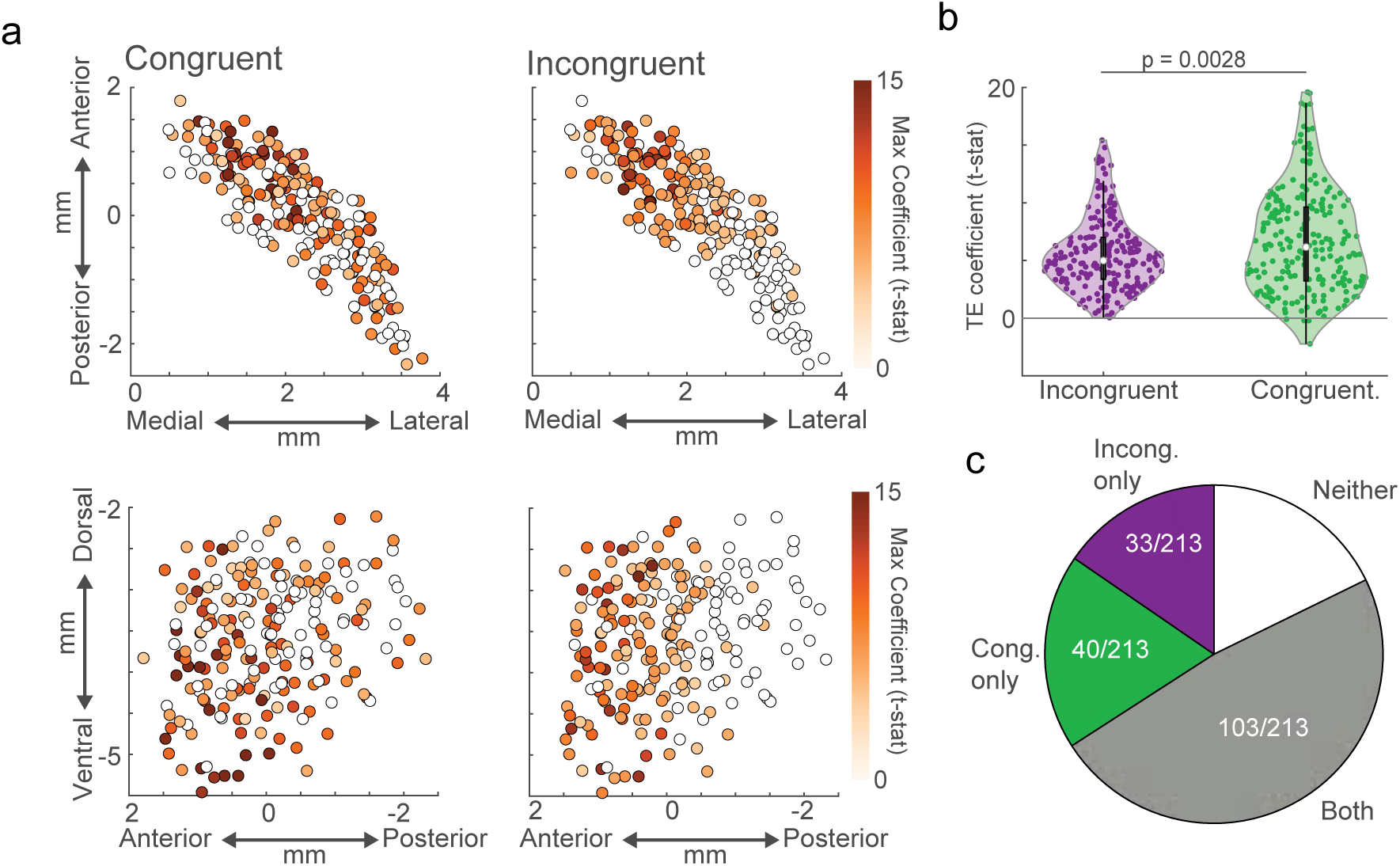
Spatial distribution of trajectory error encoding on congruent and incongruent trials independently. **a,** Horizontal (top) and sagittal (bottom) maps of max trajectory error coefficients (t-stats) for congruent (left) and incongruent (right) trials only for each fiber (circle). Open circles, not statistically significant (n = 213 fibers, 6 mice; t-test on model coefficients, p<0.05 for 3 timepoints in a row within 1s cue window, Bonferroni corrected). **b,** Violin plots of the maximum trajectory error coefficients calculated for congruent and incongruent trials separately for all fibers and mice (two-tailed shuffle test, p = 0.0028). **c,** Proportion of fibers with significant trajectory error coefficients for congruent trials only (green) incongruent trials only (purple), both (grey), or neither (white).

**Extended Data Figure 6:**
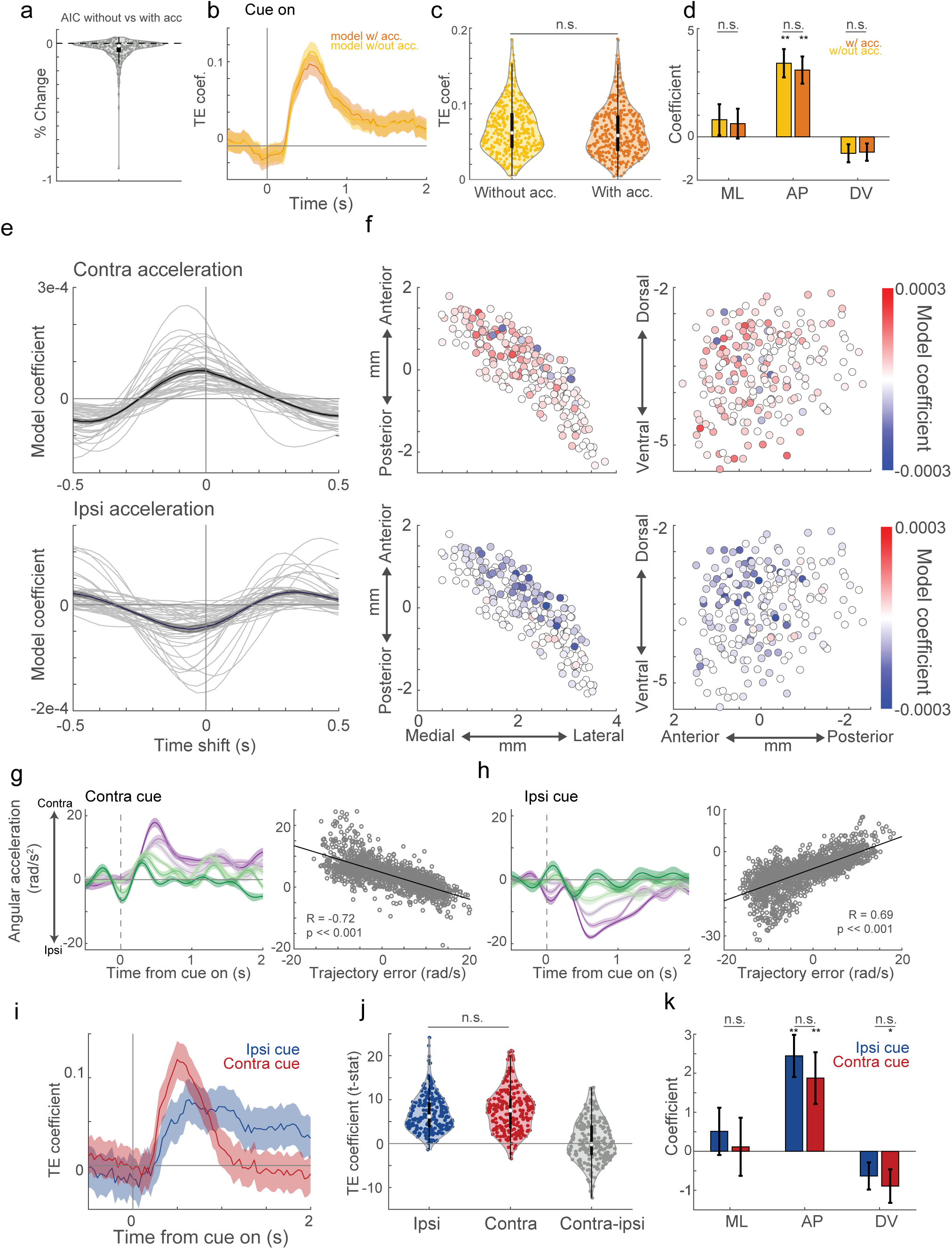
Relationships of dopamine signaling with acceleration cannot account for trajectory error encoding. **a,** Violin plots for all fibers and mice of the percent change in Akaike information criterion (AIC) from a reduced trajectory error model without acceleration variables to the full model including acceleration variables. Negative AIC changes indicate improved model fit (n = 213 fibers, 6 mice). **b,** Trajectory error coefficients for the full model (orange) and a reduced model excluding acceleration variables (yellow) for an example fiber. Shaded regions, 95% confidence interval. **c,** Violin plots of the maximum trajectory error coefficients for models with and without acceleration variables included for all fibers. (two-tailed shuffle test, n.s. p = 0.1342). **d,** Spatial coefficients for trajectory error in a model with (orange) and without (yellow) acceleration variables. Error bars, mean +/- S.E.M. (t-test on model coefficients, ANOVA on interaction *p<0.05, **p<0.01, n.s. not significant). **e,** Contralateral and ipsilateral acceleration coefficients during running in directions contralateral (top) and ipsilateral (bottom) to the implant in a model describing ΔF/F as a function of continuous acceleration at different time lags during the intertrial interval (see Methods). Acceleration was calculated from the absolute value of the angular velocity for contralateral and ipsilateral running periods independently. Each light colored line is a single fiber from one mouse, and the black line is the average across fibers. Dark grey shaded region, mean +/- S.E.M. across fibers (n = 40 fibers). **f,** Horizontal (left) and sagittal (right) maps of acceleration coefficients at zero lag during contralateral (top), and ipsilateral (bottom) running periods for each fiber (circle). Open circles are not statistically significant at zero lag (t-test on model coefficient, p<0.05). **g,** Left, average angular acceleration (+ congruent, - incongruent) aligned to cue onset on congruent (green) and incongruent (purple) contralateral cue trials, split into thirds by magnitude (dark to light colors indicates fast to slow) for one mouse (Shaded region, mean +/- S.E.M.). Right, scatterplot of average angular acceleration vs trajectory error from 0.5-1s after cue onset, for each trial across mice. Line indicates the line of best fit. Inset text indicates Pearson’s correlation coefficient and p-value (n = 1623 trials, 6 mice). **h,** Same as c, but for ipsilateral cue trials. (n = 1577 trials, 6 mice). **i,** Trajectory error coefficients aligned to cue onset for contralateral (red) and ipsilateral (blue) cue trials for one example fiber. Shaded region, 95% confidence interval. **j,** Violin plot of maximum trajectory error coefficient t-stats for contralateral cue trials, ipsilateral cue trials, and their difference (grey) across fibers and mice. (two-tailed shuffle test, n.s. p = 0.1). **k,** Spatial coefficients for trajectory error coefficients on contralateral and ipsilateral trials. (t-test on model coefficients, ANOVA on interaction *p<0.05, **p<0.01, n.s. not significant). Error bars, mean +/- S.E.M.

**Extended Data Figure 7:**
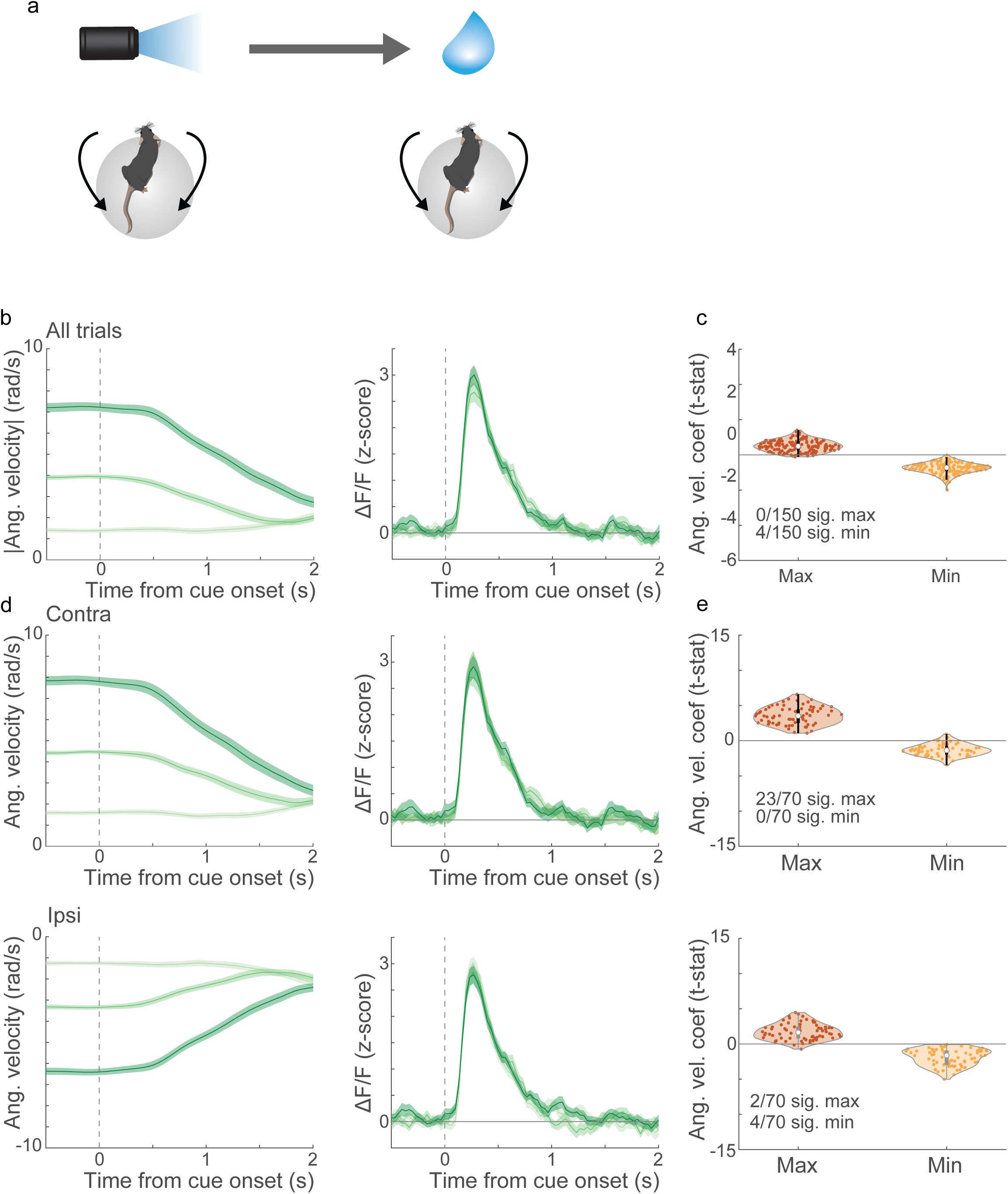
Cue-evoked dopamine release does not scale with pre-cue velocity magnitude or direction in a non-directional delay conditioning task. **a,** Schematic of non-directional delay conditioning task. Head-fixed mice running on a floating ball were presented with a fixed light cue followed by a water reward after 3 seconds, regardless of velocity. **b,** Average absolute value of angular velocity (left) and ΔF/F for a single fiber aligned to cue onset (right) across trials split into thirds based on the magnitude within 0.3 s before cue onset (dark to light colors indicate fast to slow trials). **c,** Violin plot showing the distribution of the maximum and minimum angular velocity coefficient across all fibers. Inset text, proportion of fibers significant (t-test on model coefficients, p<0.05 for 3 timepoints in a row within 1s cue window, Bonferroni corrected, n = 150 fibers, 4 mice). **d,** Same as b but split by direction (contralateral or ipsilateral relative to the implant) of the angular velocity at cue onset. **e,** Same as c, but split by direction (contralateral and ipsilateral relative to the implant) of the angular velocity at cue onset (n = 70 fibers, 2 mice). Shaded regions in all plots are mean +/- S.E.M.

**Extended Data Figure 8:**
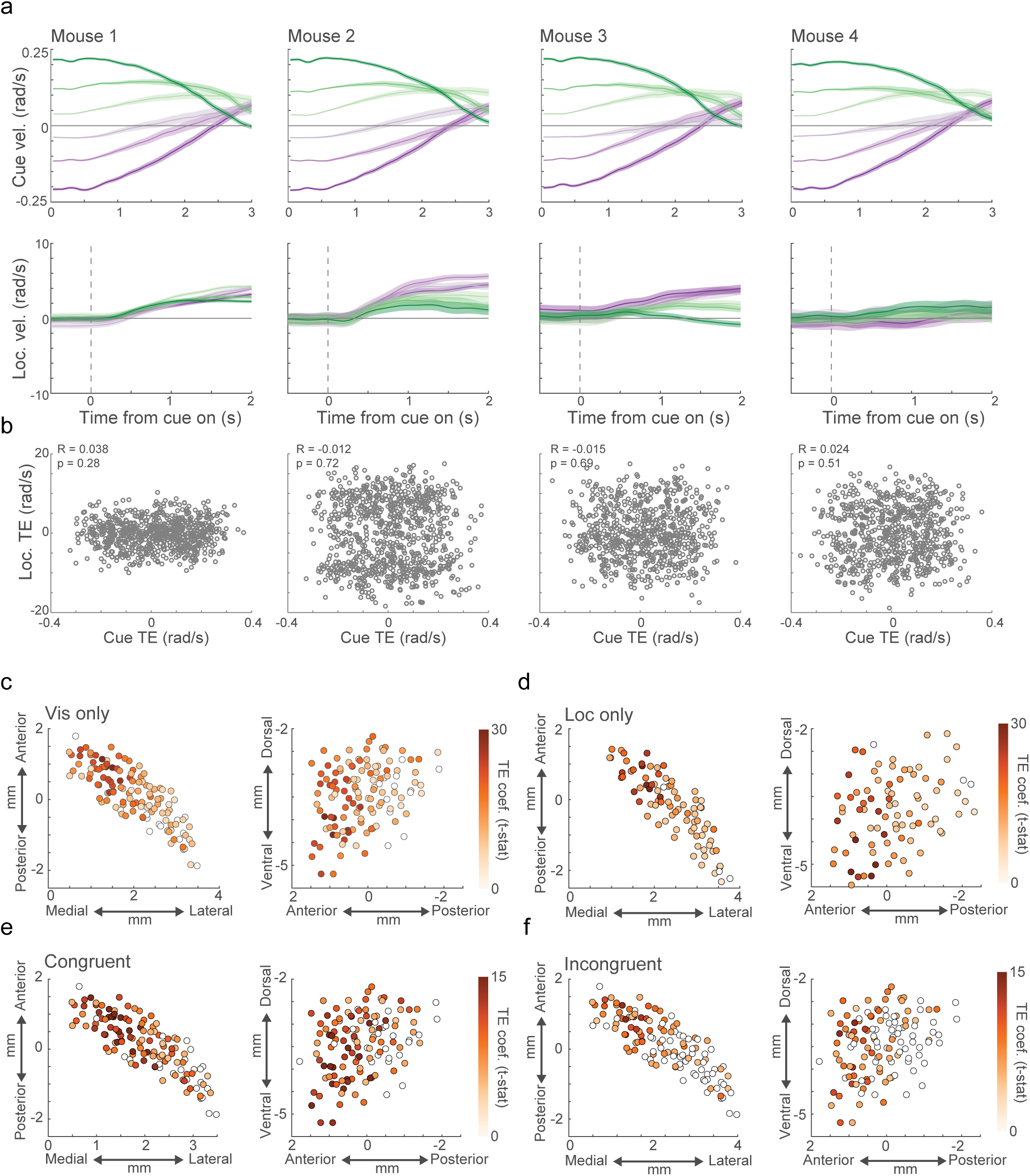
Behavior and trajectory error encoding for individual mice on the loc- and vis-only tasks. **a,** Top, average congruence-signed cue angular velocity (+ congruent, - incongruent) aligned to cue onsets in the vis-only task on congruent (green) and incongruent (purple) trials, split into thirds by magnitude for all individual mice. Bottom, average congruence-signed mouse angular velocities for the same trial groups as top. Shaded regions in all plots are mean +/- S.E.M. **b,** Scatterplots of angular velocity signed based on the mouse locomotion direction congruence vs the angular velocity signed by the cue direction congruence for all trials in the vis-only task. Inset text indicates Pearson’s correlation coefficients and associated p-values (n = 785, 901, 752, 742 trials, from left to right). **c,** Horizontal (left) and sagittal (right) maps of the maximum cue-based TE coefficients during the vis-only task for each fiber (circle). Open circles are not statistically significant (t-test on model coefficients, p<0.05 for 3 timepoints in a row within 1s cue window, Bonferroni corrected). **d,** Same as c for the loc-only task. **e,** Same as c, but for congruent (left) and incongruent (right) trials separately during the vis-only task.

**Extended Data Figure 9:**
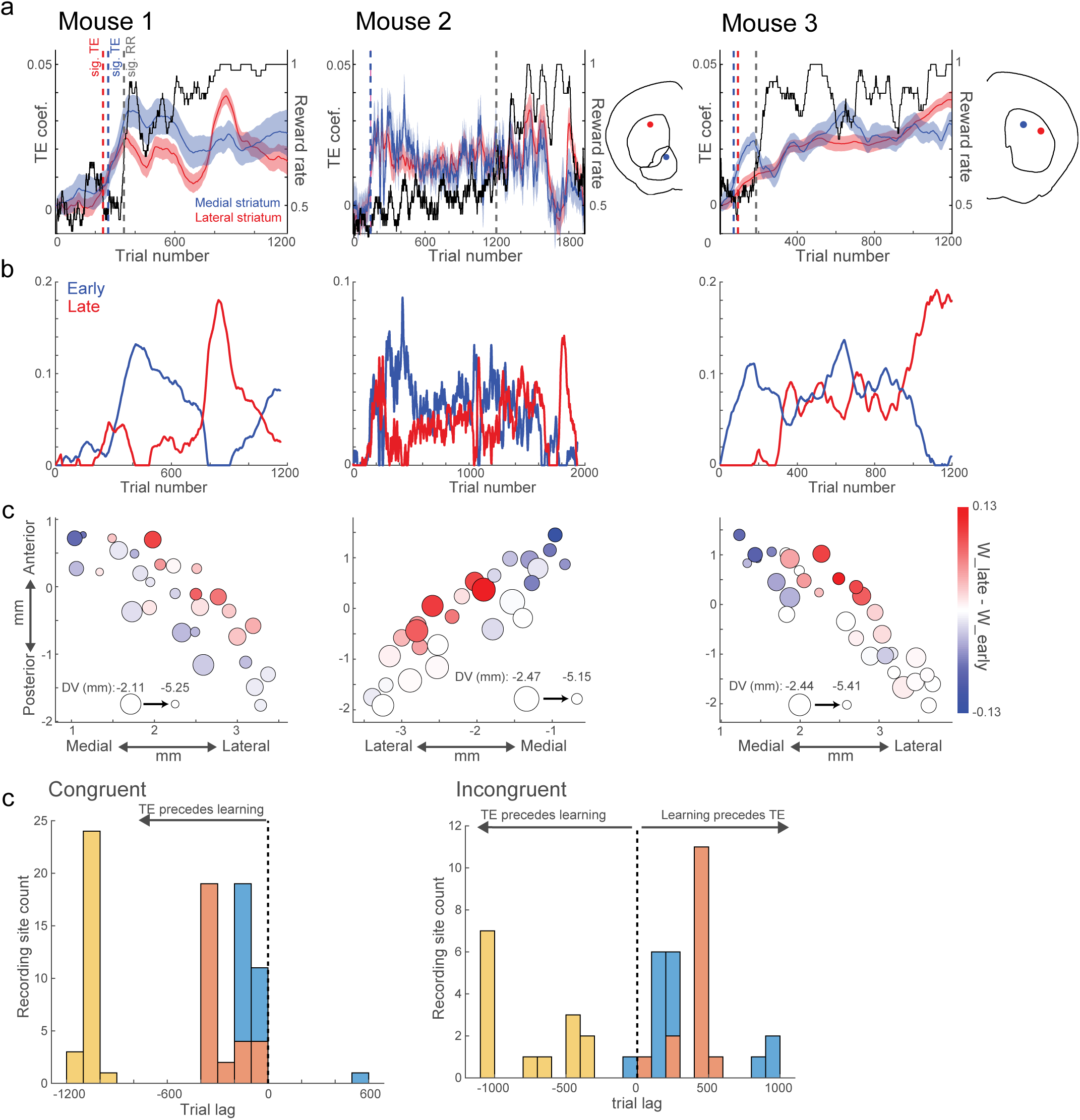
Spatial organization of the time course of TE encoding during learning is consistent across mice. **a,** Model estimates of trial by trial reward rates (black) and TE coefficients for two example fibers in each of three mice (blue, medial; red, lateral) across all trials and sessions through learning (see Methods). Dashed lines indicate the trials in which the TE coefficients and reward rates reached significance (p<0.01, shuffle test). Fiber locations indicated in coronal atlas schematics. **b,** Non-negative matrix factorization (NNMF) output factors with early (blue) and late (red) time courses for three included mice. **c,** Horizontal plane maps of the differences in the weights for the late and early components of the NNMF for each mouse. Each circle is a fiber, with the size indicating the DV position (increasing size indicates more dorsal positions). Positive values (red) indicate a stronger weight for the late component, and negative values (blue) a stronger weight for the early component. **d,** Histogram of lags between the earliest trial where congruent (left) or incongruent (right) TE and reward rate reached statistical significance for each fiber in the three included mice. Colors indicate different mice. Negative values indicate that TE becomes significant prior to reward rate.

**Extended Data Figure 10:**
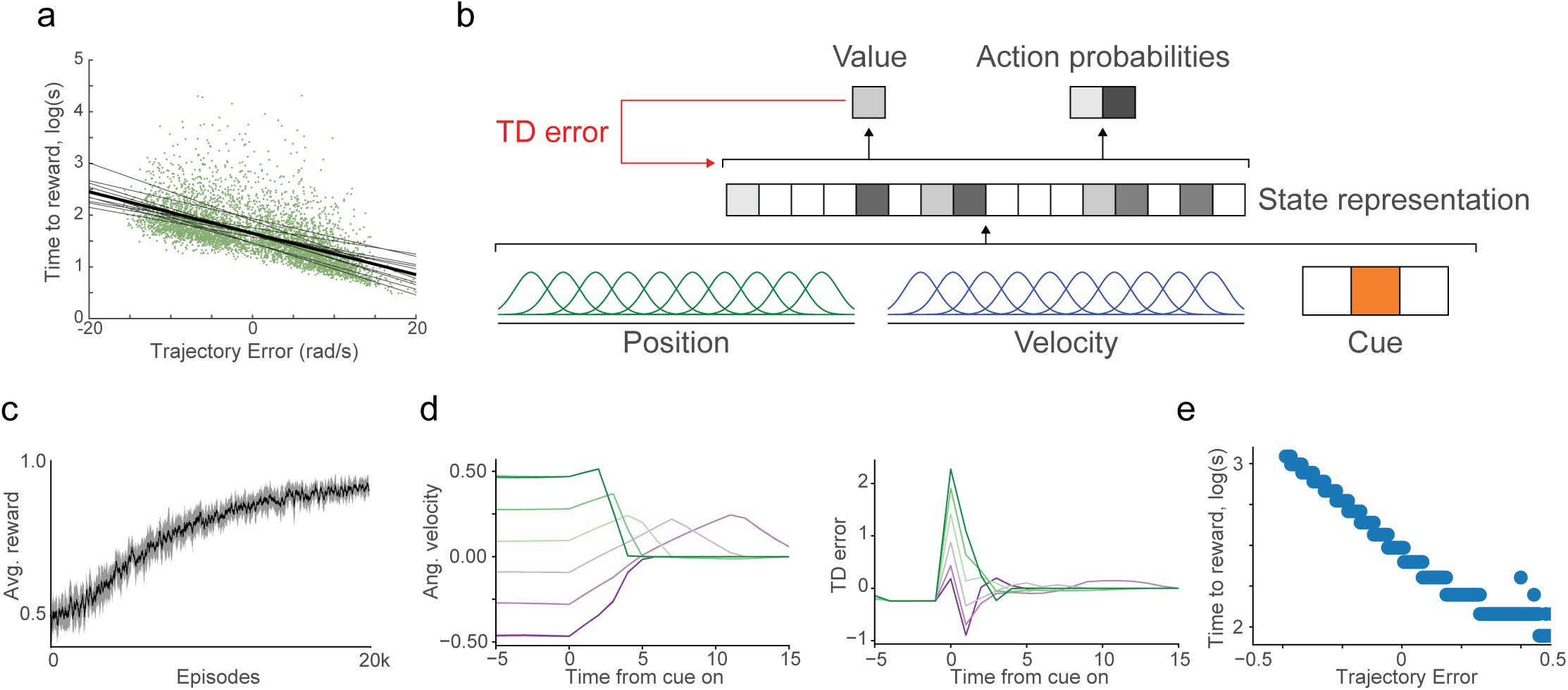
Trajectory error signaling and choice behavior can be reproduced in a reinforcement learning model. **a,** Scatter plot showing the trial-by-trial relationship between the TE and the log time to reward. Each grey line represents the least squares regression line for an individual mouse, and the black line represents the least squares regression line across mice (n = 10 mice). **b,** Schematic of a TD reward prediction error reinforcement learning algorithm. The state space was produced by a random linear mixing of a cue variable representing the presence and identity of the instructional cue and gaussian tuning curves tiling the range of positions and velocities. **c,** Average reward rate of the simulation across trials. Error bars are mean +/- S.E.M. across 10 simulations. **d,** Average velocity of the agent (left) and TD error (right) aligned to cue onset for congruent (green) and incongruent (purple) trials, split into thirds by velocity magnitude. **e,** Scatter plot showing the trial-by-trial relationship between TE and the log time to reward for one simulation.

## Notes

### Competing Interest Statement

The authors have declared no competing interest.

